# Computational Geometry for Modeling Neural Populations: from Visualization to Simulation

**DOI:** 10.1101/275412

**Authors:** Marc de Kamps, Mikkel Lepperød, Yi Ming Lai

## Abstract

The importance of a mesoscopic description level of the brain has now been well established. Rate based models are widely used, but have limitations. Recently, several extremely efficient population-level methods have been proposed that go beyond the characterization of a population in terms of a single variable. Here, we present a method for simulating neural populations based on two dimensional (2D) point spiking neuron models that defines the state of the population in terms of a density function over the neural state space. Our method differs in that we do not make the diffusion approximation, nor do we reduce the state space to a single dimension (1D). We do not hard code the neural model, but read in a grid describing its state space in the relevant simulation region. Novel models can be studied without even recompiling the code. The method is highly modular: variations of the deterministic neural dynamics and the stochastic process can be investigated independently. Currently, there is a trend to reduce complex high dimensional neuron models to 2D ones as they offer a rich dynamical repertoire that is not available in 1D, such as limit cycles. We will demonstrate that our method is ideally suited to investigate noise in such systems, replicating results obtained in the diffusion limit and generalizing them to a regime of large jumps. The joint probability density function is much more informative than 1D marginals, and we will argue that the study of 2D systems subject to noise is important complementary to 1D systems.

**Author Summary:** A group of slow, noisy and unreliable cells collectively implement our mental faculties, and how they do this is still one of the big scientific questions of our time. Mechanistic explanations of our cognitive skills, be it locomotion, object handling, language comprehension or thinking in general - whatever that may be - is still far off. A few years ago the following question was posed: Imagine that aliens would provide us with a brain-sized clump of matter, with complete freedom to sculpt realistic neuronal networks with arbitrary precision. Would we be able to build a brain? The answer appears to be no, because this technology is actually materializing, not in the form of an alien kick-start, but through steady progress in computing power, simulation methods and the emergence of databases on connectivity, neural cell types, complete with gene expression, etc. A number of groups have created brain-scale simulations, others like the Blue Brain project may not have simulated a full brain, but they included almost every single detail known about the neurons they modelled. And yet, we do not know how we reach for a glass of milk.

Mechanistic, large-scale models require simulations that bridge multiple scales. Here we present a method that allows the study of two dimensional dynamical systems subject to noise, with very little restrictions on the dynamical system or the nature of the noise process. Given that high dimensional realistic models of neurons have been reduced successfully to two dimensional dynamical systems, while retaining all essential dynamical features, we expect that this method will contribute to our understanding of the dynamics of larger brain networks without requiring the level of detail that make brute force large-scale simulations so unwieldy.

## Introduction

The population or mesoscopic level is now recognised as a very important description level for brain dynamics. Traditionally rate based models [1] have been used: models that characterize the state of a population by a single variable. There are inherent limitations to this approach, for example a poor replication of transient dynamics that is observed in simulations of spiking neurons, and various groups have proposed a population density approach. Density methods start from individual point model neurons, consider their state space, and define a density function over this space. The density function characterizes how individual neurons of a population are distributed over state space These methods have been used successfully for one dimensional point model neurons, i.e. models characterized by a single state variable, usually membrane potential. Such models, e.g. based on leaky-(LIF) or quadratic-integrate-and-fire (QIF), exponential-integrate-and-fire neurons, have a long-standing tradition in neuroscience [2–6]. Related approaches consider densities of quantities such as the time since last spike [7, 8], but here too a single variable is considered to be too coarse grained to represent the state of a population.

Recently, increased computing power and more sophisticated algorithms, e.g. [5,9–12], have made the numerical solution of time dependent density equations become tractable for one dimensional neural models. In parallel, dimensional reductions of the density have been developed, usually by expressing the density in terms of a limited set of basis functions. By studying the evolution as a time-dependent weighting of this basis the dimensionality is reduced, often resulting in sets of first order non-linear differential equations, which sometimes are interpreted as ‘rate based’ models [13–15].

The one dimensional density is very tractable: membrane potential distributions and firing rates have been shown to match spiking neuron simulations accurately, particularly in the limit of infinitely large populations, at much lower computational cost than direct spiking simulations: Cain *et al*. [16] report a speedup of two orders of magnitude compared to a direct (Monte Carlo) simulation. The problem of such one dimensional models is that they leave out details that may affect the population, such as synaptic dynamics and adaptation. Mathematically, the inclusion of variables other than just the membrane potential is no problem, but this increases the dimensionality of the state space, which negates most - but not all - computational advantages that density functions have over Monte Carlo simulation. This problem has led to considerable efforts to produce effective one dimensional methods that allow the inclusion of more realistic features of neural dynamics. Cain *et al*. have included the effects of conductances by making synaptic effects potential dependent in an otherwise standard one dimensional paradigm. Schwalger *et al*. [17] consider the distribution over the last spike times of neurons. Under a quasi-renewal approximation that the probability of a neuron firing is only dependent on the last spike time and recent population activity, they are able to model the evolution of the last spike time distribution and the population activity resulting in a system of one dimensional distributions. Both groups have modeled a large-scale spiking neuron model of a cortical column, achieving impressive agreement between Monte Carlo and density methods. Another attempt to reduce the dimensionality of the problem are moment-closure methods [18], which we will not consider here. Recently, Augustin *et al* have presented a method to include adaptation into a one-dimensional density approach [15].

There have been a number of studies of two dimensional densities [19–21]. They have made clear that analyzing the evolution of the joint probability density provides valuable insight in population dynamics, but they are not generic: it is not explicit that the method can be extended to other neural models without recoding the algorithm.

Here, we present a generic method for simulating two dimensional densities. Unlike the vast majority of studies so far, it does not start from a Fokker-Planck assumption but starts from the master equation of a point process (usually, but not necessarily) Poisson, and models the joint density distribution without dimensional reduction. We believe the method is important given the trend in theoretical neuroscience to reduce complex realistic biophysical models to effective two dimensional model neurons. Adaptive-exponential-integrate-and-fire (AdExp), Fitzhugh-Nagumo and Izhikevich neurons are examples of two dimensional model neurons that have been introduced as realistic reductions of more complex conductance based models. It is important to study these systems when subjected to noise.

The method is extremely flexible: upon the creation of a novel neural model (2D) we will be able to simulate a population subjected to synaptic input without writing a single line of new code. We require the user to present a visualization of the model in the form of the streamlines of its vector field, presented in a certain file format. Since these files can be exchanged, model exchange does not require recoding. As long as this vector field behaves reasonably - the qualification of what constitutes reasonable is a main topic of this paper - the method will be able to take it as input, and can be guaranteed to deliver sensible simulation results. The method is highly visual: it starts off with a user or stock provided visualization of a neural model, and uses computational geometry to calculate the transition matrices involved in modeling synaptic input, which is represented as a stochastic process. We will argue that with a visualization in hand one can often predict how noise will drive the system, and run a simulation to confirm these predictions. We will also show that the visualization gives a good overview of possible shapes of dynamics.

The method cannot compete in speed with effective one dimensional density methods, but holds up well compared to direct spiking neuron simulations. Since very few assumptions are used, it can be used to examine the influence of approximations made in other methods. For example, because no diffusion approximation is made, we are able to examine the influence of strong synapses, which can lead to a marked deviation from diffusion results [11, 12]. We can also model populations that are in partial synchrony.

This work captures most one dimensional population density techniques, as they are a special case of two dimensional models, in particular the method by Cain *et al*., and we also replicate results obtained in the diffusion limit as numerical solutions of Fokker-Planck equations with high precision. Although we have not tried this, theory suggests that the method should work just as well when escape noise is used [7]. With the ability to exchange neural model files, without having to recode, it is easy to check how different neural models generate dynamics in similar circuits. A software implementation of this method is available at http://miind.sf.net with a mirror repository on github https://github.com/dekamps/miind.

Since this is a methods paper, the **Material and Methods** section contains the main result, and we will present this first so that the reader may form an understanding of how the simulation results are produced. In the results section, we will show that our method works for a number of very different neural models. We will also show that strong transients, which occur in some models as a consequence of rapidly changing input, but not in others, can be understood in geometrical terms when considering the state space of the neural model.

## Materials and Methods

We will consider point model neurons with a two dimensional state space. In general such models are described by a vector field 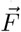, which is defined on an open subset of ℝ^2^. The equations of motion of an individual neuron are given by:

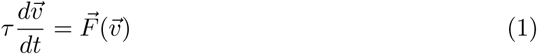

where *τ* is the membrane time constant of the neuron. We will adapt the convention that the first coordinate of 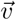 always represents a neuron’s membrane potential *v* and will refer to the second coordinate of 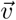 as *w*, as is conventional for the adaptation variable in the AdExp model and the recovery variable in the Fitzhugh-Nagumo model (although not in the conductance-based model). Usually boundary conditions are imposed. When a threshold potential *V*_*th*_ is present, part of *∂M*, the edge of *M*, overlaps with *V* = *V*_*th*_. This part of *∂M* is called the threshold. When a neuron state approaches the threshold from below, the state is reset, sometimes after a refractive time interval *τ*_*ref*_ during which its state is effectively undefined. The reset results in coordinate *v* being set to a reset potential *V*_*reset*_, whilst the second coordinate remains unaffected if no refractive interval or period is considered. If there is a refractive period, there are variations: sometimes the second coordinate is kept constant, sometimes further evolution according to Eq. (1) for a period of *τ*_*ref*_ is considered and the reset value of the second coordinate is taken to be the resulting value of *w*(*t*_*spike*_ + *τ*_*ref*_), where *t*_*spike*_ is the time when the neuron hits threshold. The neuron itself emits a spike upon hitting the threshold. This description fits many neuron models: e.g. adaptive-integrate-and-fire; conductance-based leaky-integrate-and-fire; Izhikevich [44], and many others.

We are interested in populations of neurons. We consider a population to be homogeneous: all neurons have the same parameters, and statistically see the same input: they are subject to input spike trains generated from the same distribution. Under those considerations one can define a density, 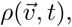 over state space for a population that is sufficiently large. 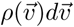 is defined as the fraction of neurons in the population whose state vector is in 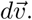 For spike trains generated by a Markov process, the evolution equation of the density obeys the differential Chapman-Kolmogorov equation:

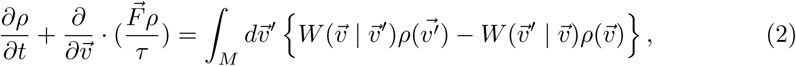

where 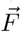 and *τ* are from the neuron model as stated in Eq. (1).

Input spikes will cause instantaneous responses in the state space of neurons. For delta synapses, for example, an input spike will cause a transition from membrane potential *V* to membrane potential *V* + *h*, where *h* is the synaptic efficacy, which may be drawn from a probability distribution *p(h).* In current based models the jump may be in the input current, and in conductance based models, studied below, the jump is in conductance, rather than membrane potential. Nonetheless, in all of these cases the input spikes cause instantaneous transitions from one point in state space to another. The right-hand side of Eq. 2 expresses that the loss of neurons in one part of state space is balanced by their reappearance in another after the jump. As a concrete example, consider input spikes generated by a Poisson point process with delta synapses:

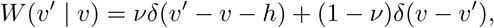

where *v* is the rate of the Poisson process and *h* is the synaptic efficacy, which for simplicity we will consider here as a single fixed value. Eq. 2 reduces to:

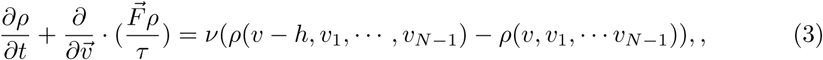

where the *v*_*i*_ are the components of 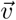.

At this stage, often a Taylor expansion is made for the right-hand side of the equation up to second order, which leads to a Fokker-Planck equation. We will not pursue this approach, instead we will point out, as observed by de Kamps [10,12] and Iyer *et al.* [11] that the method of characteristics can be used to bring Eq. 3 into a different form. Consider a line segment *l* in state space, and pick a point 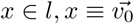 at *t* = 0. The system of ordinary differential equations Eq. 1 defines a curve that describes the evolution of point 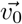 through state space. This curve is an integral curve of the vector field 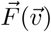 and can be found by integration. Writing this curve as 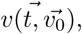 we can introduce a new coordinate system:

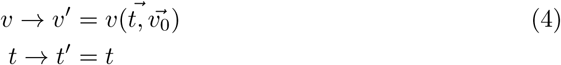

In this new coordinate system Eq. 2 becomes:

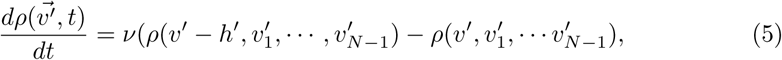

which has the form of a Poisson master equation. This implies that rather than solving the partial integro-differential equation Eq. 2, we have to solve the system of ordinary differential equations Eq. 5. This system describes mass transport from bin to bin and no longer has a dependency on the gradient of the density profile: the drift term in Eq. 2 has been transformed away. Equation 5 describes mass transport from one position to another. The distance between these positions is now immaterial and this means that arbitrary large synaptic efficacies can be handled.

The observation that for a system that co-moves with the neural dynamics all mass transport is determined by the stochastic process is important. It suggests that the right-hand side of Eq. 5 - representing the master equation of a Poisson process - can be replaced by more general forms without affecting the left-hand side of the equation that allows use of the method of characteristics. Indeed, recently we have considered a generalization to spike trains generated by non-Markov processes [23]. This generalizes the right-hand side of Eq. 2, but leaves the left-hand side unchanged, and in [23] we show explicitly that for one dimensional densities the method discussed here extends to non-Markov renewal processes. The generalization of Eq. 2 requires a convolution over the recent history of the density, using a kernel whose shape is dependent on the renewal process.

Consider a two dimensional state space with coordinates *v* and *w.* The coordinate transformation just described defines a mapping from point *x* on a line segment of initial points to a point in state space:

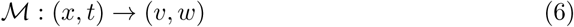

This has two implications: first, the evolution of the initial line segment *l* over a given fixed period of time defines a region of state space. The state space relevant to a simulation may have to be built from several such regions. Second, the mapping is time-dependent: Eq. 4 must be solved in a coordinate system that itself is subject to dynamics: that of the deterministic neuron. This suggests a solution consisting of two interleaved steps: one accounting for deterministic movement of neurons, and one where Eq. 5 is solved numerically. We will now describe this process in detail.

### State Space Models of Neuronal Populations

As an example, we consider a conductance based model with first order synaptic kinetics following [20]. It is given by:

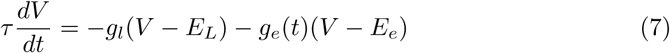

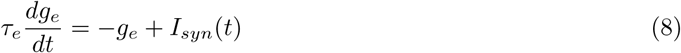

Numerical values are taken from [20], and given in Table 1. *I*_*syn*_(*t*) represents the influence of incoming spikes on the neurons. A conventional representation of such a model is given by a vector field, see Fig. 1.

**Table 1.** Constants taken from Apfaltrer et al. (2006), Appendix B.

|  |  |  |
| --- | --- | --- |
| membrane time constant | $\tau_m$ | 20 ms |
| reset potential | $E_r$ | -65 mV |
| reversal potential | $E_{rev}$ | -65 mV |
| equilibrium potential $e$ | $E_e$ | 0 V |
| synaptic time constant $e$ | $\tau_s$ | 5 mV |
| threshold potential | $V_{th}$ | -55 mV |

**Fig. 1.**
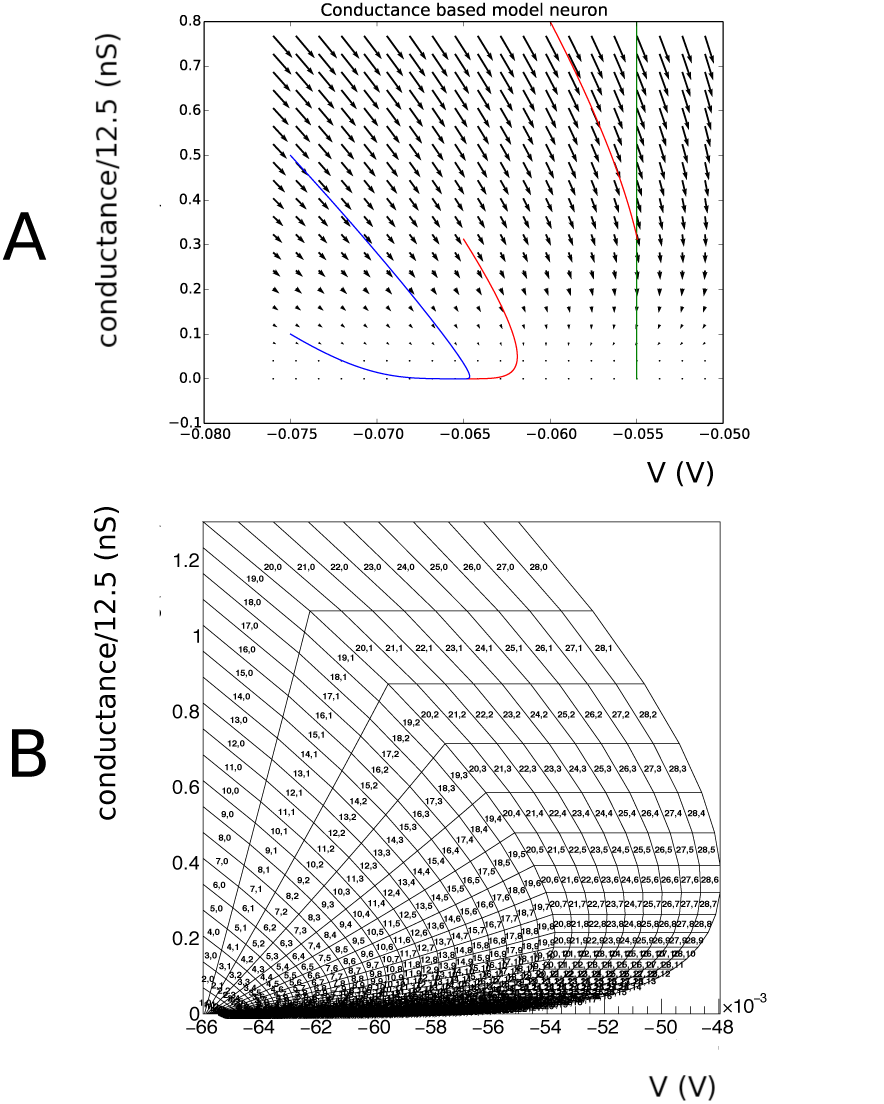
A: Vector field for a conductance based model, along with a few integral curves. At very low conductance, there is a drive towards equilibrium regardless of the initial point. At higher conductance values the drive is dominated by a trend towards the equilibrium potential of the excitatory synapse (0 mV). The blue integral curves demonstrate this. The red integral curve represents a neuron that hits the threshold potential (−55 mV), and subsequently undergoes a reset to the reset potential (−65 mV). This neuron will emit a spike. After reset, it will not hit threshold again and eventually asymptotes to equilibrium potential (−60 mV). B An example grid for the conductance based model. The grid is built from strips. Strip numbers are arbitrary, as long as they are unique, but it is convenient to number them in order of creation. By construction, cell numbers within a strip are ordered by the dynamics: neurons that are in cell number *j* of strip *i* at time *t* are in cell number *j* + 1 mod *n*_*j*_ of strip *j* at time *t* + Δ*t*, where *n*_*j*_ is the number of cells in that strip.

• A number of initial points are taken:

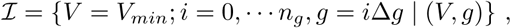

for given fixed *V*_*min*_, *n*_*g*_, Δ*g*

Consider a two dimensional dynamical system defined by a vector field. A point in state space will be represented by a two dimensional vector 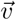. A grid is constructed from strips. As mentioned previously, usually one dimension is a membrane potential, and we will denote coordinates in this dimension by a small letter *v*. The second dimension can be used to represent parameters such as adaptation, conductance, and will be represented by *w.* A strip is constructed by choosing two neighbouring points in state space, e.g. 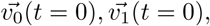 and integrating the vector field for a time *T* that is assumed to be an integer multiple of a period of time *Δt*, which we assume to be a defining characteristic of the grid. Let *T* = *nΔt*, then two discretized neighbouring characteristics

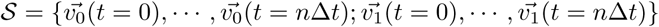

define a strip. Within a strip, the set of points

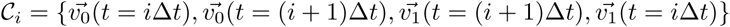

defines a cell, which is quadrilateral in shape. The quadrilateral should be *simple*, but not necessarily *convex* (Fig. 2 A). We reject cells with less than a certain area. As we will see in concrete examples, boundaries in state space are approached through areas of vanishing measure. The area cut tends to remove complex cells, and we will reject them in general. An example of a grid generated by this procedure is given in Fig. 3.

**Fig. 2.**
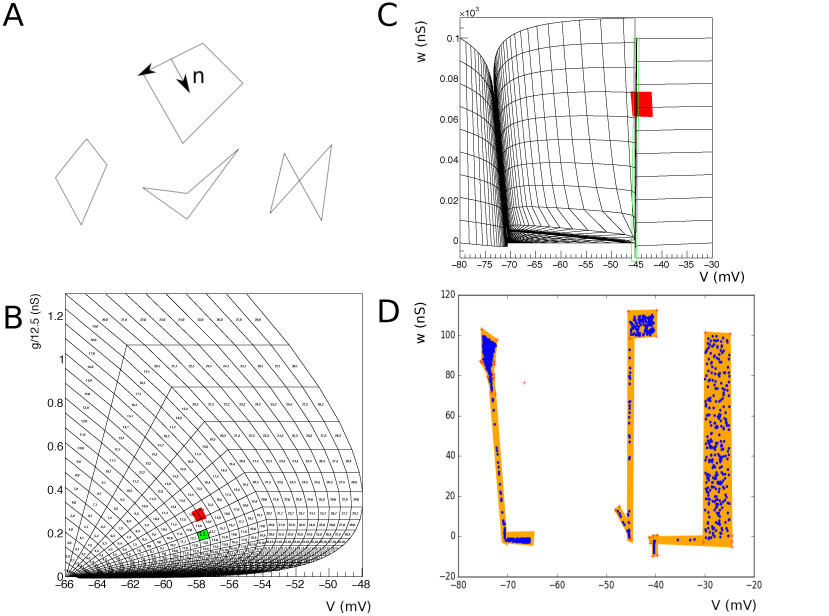
A: As a result of the integration procedure simple quadrilaterals (left, middle) should emerge, which are usually convex (left), except near stationary points or limit cycles where concave quadrilaterals (middle) can be formed. Complex, i.e. self-intersecting, quadrilaterals can occur around strong non linearities, for example the crossing of nullclines. These definitions hold for any polygon. B: The problem of defining the Master equation: we can easily calculate how much mass per unit time leaves a given bin *(i,j).* This mass will reappear at a position *h* away from the original bin, where *h* is the synaptic efficacy. In the figure bin (13,7) is translated along vector (0,0.1). This corresponds to neurons that have received an input spike, and therefore are experiencing a jump in conductance. Most neurons that are in bin (13,7), will end up in bin (13,5) and (14,4), with some in bin (13,6) and (14,6). So *C*_(0., 0.1)_ (13, 7) = {(13, 5), (14, 5), (13, 6), (14, 6)}. C: Some events will end up outside of the grid after translation. D: Fiducial quadrilaterals can be used to test where they have gone missing, and where is the best place to reassign them to the grid.

**Fig. 3.**
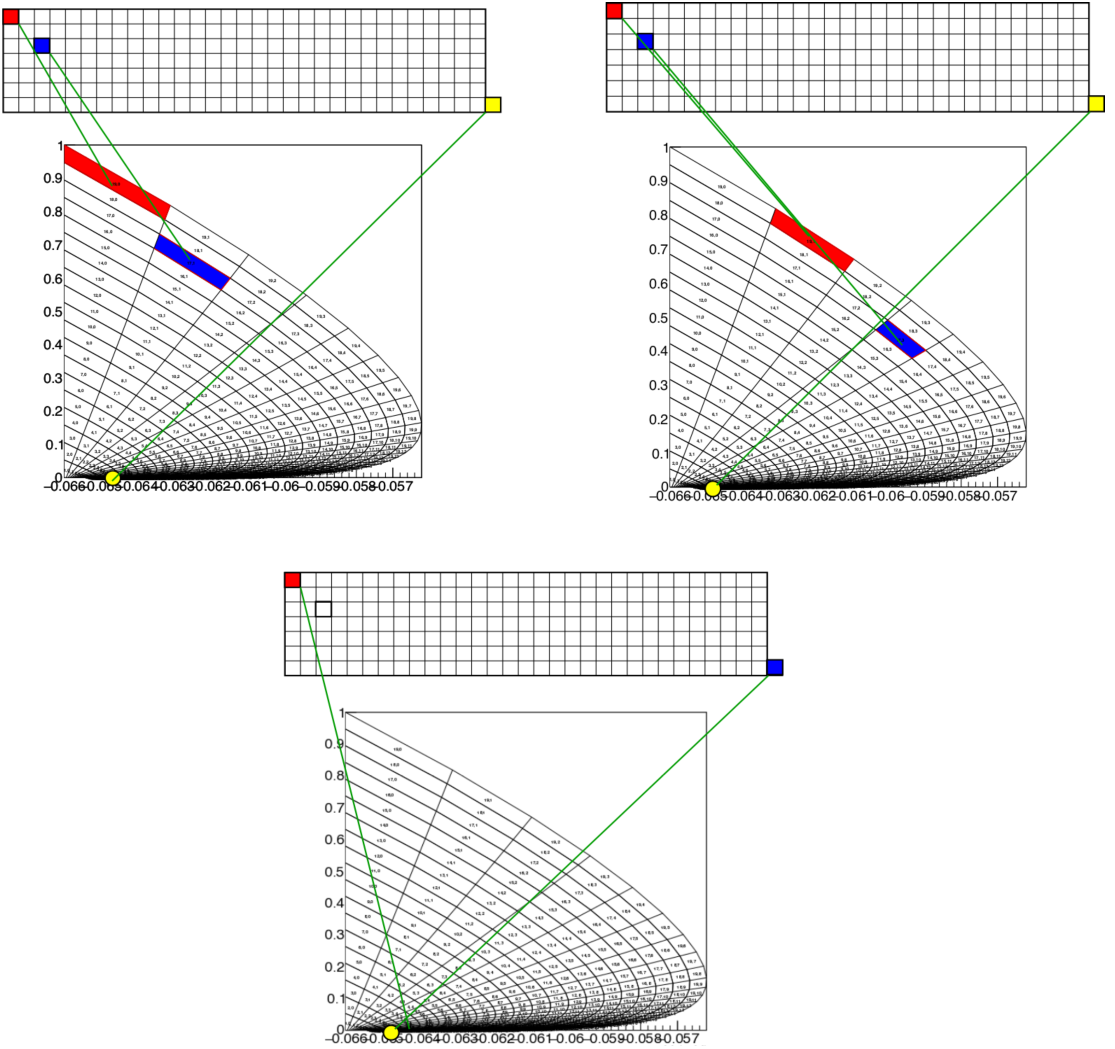
Probability mass is maintained in a mass array. In general, mass does not move, except when the mass has moved beyond the end of a strip. The relationship between the mass array and the mesh is updated with each time step, resulting in the apparent motion of probability mass through the mesh (top left and top right). At the end of each simulation step, probability mass is removed from each first bin of the strip, and added to a special quadrilateral (bottom): the reversal bin. Mass does not move from here, and only synaptic input can cause mass to leave this bin.

Strip numbers are arbitrary, as long as they are unique, but it is convenient to number them in order of creation. In the remainder of the paper, we will assume that strip numbers created by the integration procedure start at 1, and are consecutive, so that the numbers *i* ∊ {1, …, *N*_*strip*_} with *N*_*strip*_ the number of strips, each identify a unique strip. Strip no. 0 is reserved for stationary points. There may be 0 or more cells in strip 0. The number of cells in strip *i* denoted by *n*_*cell*_*(i)*. We refer to the tuple *(i,j)*, with *i* the strip number and *j* the cell number, as the *coordinates* of the bin in the grid. *N*_*cells*_ is the total number of cells in the grid.

For all strips *i* (*i* > 0 by construction), cell numbers within a strip are ordered by the dynamics: neurons that are in cell number *j* of strip *i* at time *t* are in cell number *j* + 1 mod *n*_*j*_ of strip *j* at time *t + Δt*, where *n*_*j*_ is the number of cells in that strip.

Neurons that are in a cell in strip no. 0 are assumed to be stationary and do not move through the strip. Examples of cells in this strip are reversal bins. The handling of stationary bins will be discussed below.

### Representing a Density Profile

A simulation progresses in multiple steps of Δ*t*, so the current simulation time *t*_*sim*_ is specified by an integer *k*, defined by:

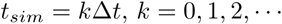

The density profile can be represented in an array *ℳ* of length *N*_*cells*_. Each element of this array is associated with the grid as follows. Let *c*_*cell*_(0) ≡ 0 and for 0 < *i* ≤ *N*_*strip*_ let *c*_*cell*_*(i)* ≡ *c*_*cell*_ (*i* - 1) + *n*_*cell*_(*i* - 1), so *c*_*cell*_ *(i)* represents the total number of cells in all strips up to strip *i.* Now define the index function *I*:

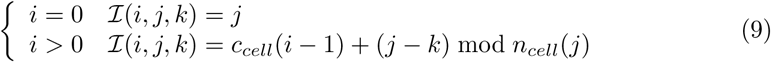

This is a time dependent mapping: its effect is a forward motion of probability mass with each forward time step. We will refer to the updating of the mapping by incrementing *k* as a *mass rotation as* probability mass that reaches the end of a strip, will reappear at the beginning of the strip at the next time step. This effect is almost always undesirable as it would effect a jump wise displacement of probability mass. In most models this can be prevented by removing the probability mass from the beginning of each strip and setting the content of this bin to 0, and adding the removed mass to a another bin. A typical example arises in the case of integrate-and-fire models. Here, there is usually a reversal point. Such a point can be emulated by creating a small quadrilateral, and making this cell number 0 in strip number 0.

The procedure of mapping probability mass from the beginning of a strip to special bins in state space is called a *reversal mapping*. It consists of a list of coordinate pairs. The first coordinate labels the bin where probability will be removed, the second coordinate labels the bin where the probability will reappear. The concept of reversal mapping extends to other neural models - we will consider adaptive-exponential-integrate-and-fire (AdExp), Fitzhugh-Nagumo, and quadratic-integrate-and-fire neurons. All of these models need a prescription for what happens with the probability mass after reaching the end of a strip, and we will refer to this as the reversal mapping, even if the model does not really have a reversal bin, to contrast it from the *threshold mapping*. Although handling a threshold is similar, interaction with synaptic input means that the mapping requires extra precautions. We will discuss this in the section below.

The whole process of advancing probability through a grid by means of updating a relationship with a grid is illustrated in Fig. 2. Up to this point we have only referred to probability mass. If a density representation is desired, one can calculate the density by:

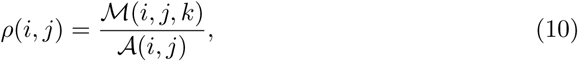

where *A(i, j)* is the area of quadrilateral *(i,j)*, and *M(i,j, k)* is the probability mass present in the quadrilateral *(i,j)* at simulation time *kΔt*. We note that this procedure implements a complete numerical solution for the advective part of Eq. 2.

### Handling Synaptic Input

We will assume that individual neurons will receive Poisson spike trains with a known rate for a known synaptic distribution of the post synaptic population. Without loss of generality we will limit the exposition to a single fixed synaptic efficacy; continuous distributions can be sampled by generating several matrices, one for each synaptic efficacy, and adding them together. Adding the individual matrices, which are band matrices, and very sparse, results in another band matrix, still sparse, albeit with a slightly broader band. Overall run times are hardly affected unless really broad synaptic distributions are sampled.

A connection between two populations will be defined by the tuple (*N*_*con*_, *h, τ*_*delay*_). Here *N*_*con*_ is the number of connections from presynaptic neurons onto a representative neuron in the receiving population, *τ*_*delay*_ the delay in the transmission of presynaptic spikes and *h* the synaptic efficacy. The firing *v* rate is either given, or inferred from the state of the presynaptic population, but in both cases assumed to be known. For the population these assumptions lead to a Master equation:

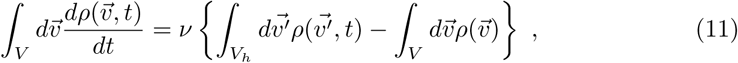

where *V* is an area of state space and *V*_*h*_ the same area, translated by an amount *h* in dimension *i.* It is dependent on the neuronal model in which variable the jump takes place. In AdExp the jump is in membrane potential, in conductance based models it is in the conductance variable. Here, we will discuss the problem using conductance based neurons as an example, but the methodology applies to any model.

Eq. (11) determines the right hand side of Eq. (2), and the stage is set for numerical solution. The left hand side of Eq. (2) describes the advective part, and is purely determined by the neuron model, which ultimately determines the grid. We already have described the movement of probability mass due to advection during a time step Δ*t*, and need to complete this by implementing a numerical solution for Eq. (11).

Eq. (11) describes the transfer of probability mass from one region of state space to another. We will assume that the grid we use for the model of advection is sufficiently fine, so that the density within a single bin can be considered to be constant, and choose area *V* in Eq. (11) to coincide with our grid bins. We approximate (11) by:

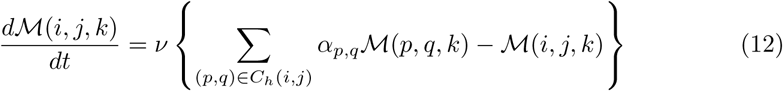

The bin *(i,j)* translated by a distance *h* will cover a number of other bins of the grid. Let (*p, q)* be a bin partly covered by the translated bin *(i,j)* and let *α*_*p, q*_ be the fraction of the surface area of the translated bin that covers bin *(p, q).* (By construction 0 < *α*_*p,q*_ ≤ 1.) The set *C*_*h*_(*i,j*) is defined as the set of tuples (*p, q*), for all such bins, i.e. those bins that are covered by translated bin *(i,j)* (and no others). We will refer to *C*_*h*_(*i,j*) as the *displacement set*. Usually, the displacement is in one dimension only, where this is not the case we will write 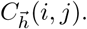 The problem of determining *C*_*h*_(*i,j*) is one of computational geometry that can be solved before simulation starts. It is illustrated in Fig. 2 B, where the grid of the conductance based model is shown.

This problem is easily stated but hard to solve efficiently. Conceptually, a Monte Carlo approach is simplest, and since the computation can be done offline - before simulation starts - this approach is preferable. It is straightforward for a given bin of the grid *(i,j)* to generate random points that are contained within its quadrilateral. Assume these points are translated by a vector 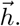 It is now a matter of determining in which bin a translated point falls. In order to achieve this the grid is stored as a list of cells. Each cell, being a quadrilateral, is represented by a list of four coordinates. During construction of the grid, vertices of a cell are stored in counter clockwise order.

When a quadrilateral is convex, and the vertices are stored in counter clockwise order, the × operator defined by:

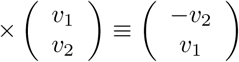

results in an “inward” pointing normal 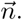 If the position vector of a point has a positive scalar product with the ‘inward’ normal of all four line segments that define the quadrilateral the point is inside, otherwise it is outside. These half line tests are cheap and easy to implement. If the quadrilateral is not convex, but simple, it can be split into two triangles which are convex.

We perform linear search to find a grid cell that contains the translated point, or to conclude there is no such cell. Better efficiency can be obtained with k-d trees, but we have found the generation of translation matrices not to be a bottleneck in our workflow, and linear search allows straightforward brute force parallelization. At most one cell will contain the translated point. For now, we will assume that the translated point will be inside a given bin (*p, q*). Later, for concrete neuron models we will discuss specific ways of handling transitions falling outside the grid. If bin (*p, q*) is not represented in 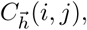 an entry for it will be added to it. The process is then repeated, in total *N*_*point*_ times. For each cell *(p,q)* represented in 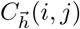> a count *n*(_*p,q*_) is maintained and *α*_*p,q*_ is estimated by:

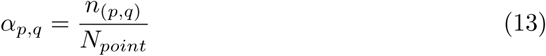

Equation 12 is of the form

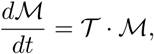

where *T* is called the transition matrix. The displacement set determines the transition matrix.

Here, we have described a Monte Carlo strategy that uses serial search to determine the set 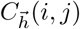 and consequently the constants *α*_*p,q*_ for bins *(p,q)* in that set. With these constants determined, it is a straightforward matter to solve Eq. 12 numerically.

The main algorithm now consists of three steps: updating the index relationship Eq. 9, which constitutes the movement of probability mass through the grid during a time interval Δ*t*; implementing the reversal mapping; solving Eq. 12 during Δ*t*. The order of these steps matters. Implementing the reversal bin after the master equation may lead to removing probability mass from the beginning of the strip that should have been mapped to a reversal bin.

### Handling a Threshold

Many neuron models incorporate a threshold of some sort. For example, in the original conductance based model by [20], a threshold of-55 mV is applied. This corresponds to a vertical boundary in the *(V,g)* plane (see Fig. 1). Neurons that hit this threshold from lower potentials generate a spike and are taken out of the system. After a period *τ*_*ref*_, they are reintroduced at (*V*_*reset*_, *g(t*_*spike*_ + *τ*_*ref*_)), where *t*_*spike*_ is the time when the neuron hits the threshold, and *g*(*t*_*spike*_) is the conductance value the neuron had at the time of hitting the threshold. In this model, following [20], it is assumed that the conductance variable continues to evolve according to Eq. 8, without being affected by the spike.

We handle this as follows. For each strip it is determined which cells contain the threshold boundary, i.e. at least one vertex lies below the threshold potential and at least one lies on or above the threshold potential. The set of all such cells is called the *threshold set.* In a similar way a *reset* set is constructed, the set of cells that contain the reset potential. In the simplest case, for each cell in the threshold set the cell in the reset set is identified that is closest in *w* to that of a threshold cell. The threshold cell is then mapped to the corresponding reset cell and the set of all such mappings is called the reset mapping.

Sometimes, the value of *w* is adapted after a neuron spikes. In the AdExp model, for example, *w* → *w* + *b* after a spike. In this case, we translate each cell in the reset set in direction (0,*w*), and calculate its displacement set, just as we did for the transition matrix. The reset mapping is then not implemented between the threshold cell and the original reset cell, but to the displacement set of that reset cell. We do this for all threshold cells and thus arrive at a slightly more complex reset mapping.

Due to the irregularity of the grid, it may happen that some transitions of the Master equation are into cells that are above the threshold potential. This will lead to stray probability above threshold, if not corrected. We correct for this during the generation of the transition matrix. If during event generation a point ends up above threshold after translation, we look for the closest threshold cell for this point. The event is then attributed to that threshold cell, and not the stray cell above threshold. In this way transitions from below or on threshold to cells above threshold are explicitly ruled out.

The reset mapping must be carried out immediately after the solution of the master equation, before the next update of the index function.

### Gaps in State Space

All grids are finite. For that reason alone the Monte Carlo procedure described above will result in translated points that cannot be attributed to any cell. Those events are lost and will lead to unbalanced transitions: mass will flow out of bins near the edge, but will not reappear anywhere else in the system and there is a possibility that mass evaporates from the system. This problem does not occur just at the edges, but also in the vicinity of stationary points. We will see that some dynamical systems display strong non linearities that will make it impossible to cover state space densely. The ability to deal with such gaps in state space is the most important technical challenge for this method.

In Fig. 2 we show how to handle these gaps. Figure 2 B shows that a cell which is translated by 5 mV can fall across a small cleft not part of the grid. We cover this gap by a quadrilateral (in green): a *fiducial* cell. An event that is not within the grid, but inside this quadrilateral needs to assigned to a mesh cell, otherwise the transition matrix will not conserve probability mass. It is straightforward to maintain a list of grid cells that have at least one vertex in the fiducial bin. We assign the event to the grid cell that is closest along the projection in the jump direction.

Figure 2 D shows the total number of events lost in the generation of transition matrix corresponding to a jump of 5 mV, thereby revealing gaps in state space. The orange quadrilaterals are the fiducial bins. After reassignments all events fall inside the grid and probability will be balanced.

### Marginal Distributions

It is straightforward to calculate marginal distributions. Again, we use Monte Carlo simulation to generate points inside a given quadrilateral (*p, q).* We then histogram these points in *v* and *w.* For each bin *i* in the *v* histogram, we can now estimate a matrix element α_*(p,q),i*_ by dividing the number of points in bin *i* by the total number of points that were generated. For a given distribution, one can now multiply the total mass in bin (*p, q*) by *α*_(*p,q),i*_ to find how much of this mass should be allocated to bin *i*. If one does this for every cell (*p, q)* in the grid, one will find the distribution of mass over the marginal histogram, and can calculate the marginal density from this.

## Results/Discussion

We present a succession of population simulations of four neuronal models. A neuron with a single excitatory conductance has a simple state space, and its simulation provides few problems. It is a familiar model and therefore a good one to introduce and demonstrate the formalism. We then move to one dimensional results and replicate some familiar results for LIF and QIF neurons: density profiles, transient firing rates and gain curves. This allows us to quantitatively examine some of the strengths and weaknesses of the method. We then discuss two models that show a progression of difficulty in covering state space: AdExp and Fitzhugh-Nagumo. Although a methods paper, we feel that nonetheless we can infer a number of general principles that run as a common thread through our use cases, and we present them here.

- We obtain a general method for simulating populations of spiking point model neurons with a one or two dimensional state space, subject to Poisson spike trains. When restricted to one dimension, the method is equivalent to that published by de Kamps (2013) and Iyer *et al.* (2013) and is very efficient, as the work of Cain *et al.* demonstrates. The method is able to replicate earlier work on 2D models, but is more general, as first, it is able to accept novel models in the form of a grid file and therefore does not require source code changes when a new model is considered, and second, does not rely on the diffusion approximation, but allows a variety of stochastic processes to be considered. The method is most efficient for synaptic efficacies and firing rates commensurate with what is found in the brain, but can be pushed to reproduce diffusion results, although dedicated numerical strategies for solving the ensuing 2D Fokker-Planck equations will be more efficient. Nonetheless, the possibility to study the diffusion limit as a special case is a useful property of the method.
- The method is insensitive to the gradient density, and will accurately model delta synapses and handle discontinuities of the density profile, and is able to model populations that are in partial synchrony, allowing the modelling of the decorrelation process itself.
- The neural model will be presented in a file representation of a state space diagram. For some models it is hard to cover state space completely due to singularities, for example when approaching nullclines. Such parts of state space are effectively forbidden for endogenous deterministic neural dynamics, but noise may place events there, moving neurons outside state space. We find there are two cases where this happens: first, on the approach of one of the nullclines the system approaches a stable equilibrium or a limit cycle. The system does not contain enough information in one of the two dimensions and the grid cannot be meaningfully continued. We find that the motion of probability mass inside such a region can be inferred from the dynamics around it. A limit cycle, for example can be inferred from the grid closing onto it, even when we cannot extrapolate the grid directly to the limit cycle. In a similar way we can capture the motion of mass towards a stable equilibrium: when motion has stopped in one direction, but still continues in the other, we find that placing neurons that are deposited into accessible regions by synaptic input in nearby parts of state space accurately captures the overall motion of mass around these regions. Similar considerations apply around unstable regions of state space, and because we can time invert the dynamical system when constructing the grid, we find that these problems can be handled in much the same way.
- Transient responses can be understood in geometrical terms. If a boundary, either a reflecting or an absorbing one, is present in state space, the population will exhibit a strong oscillatory response (“ringing”) when the input is strong enough to push neurons towards the boundary and noise is too weak to disperse neurons before reaching it. The converse is also true: if despite the presence of a boundary, state space allows neurons a way around it, strong transients will be absent. Rate based models based on first order differential equations on using a gain function will model these transients incorrectly, or not at all.
- The method can describe the version of Tsodyks-Makram synapses used by Vasilaki and Giugliano [22] in a model of network formation.
- By far the most challenging grid to make was that of a Fitzhugh-Nagumo neuron, because the approach to the limit cycle in part also implies an approach to the nullcines of the system, leading to a loss of information in one dimension. Where the nullclines cross this problem is exacerbated. We find that we have to imply the limit cycle: we define the grid in the approach to the limit cycle and infer the deterministic dynamics in an area around the limit cycle from the surrounding grid cells.

### Conductance Based Neurons

We consider neurons with a single excitatory synapse as given by Eq. (8). In Fig. 4 we present first the simulation of a jump response: a group of neurons is at rest at time *t = 0* and all neurons are at (*V = −65 mV, g = 0*). From *t* = 0 onward the neurons will receive Poisson distributed input spike trains with a rate of 1000 Hz. A neuron that receives an input spike will undergo an instantaneous state transition and move up in conductance space. Until it receives a further input spike it will start to move through state space under its endogenous neural dynamics: the neuron will depolarize and simultaneously reduce its conductance. The process was described in Sec. **Materials and Methods: State Space Models of Neuronal Populations**.

**Fig. 4.**
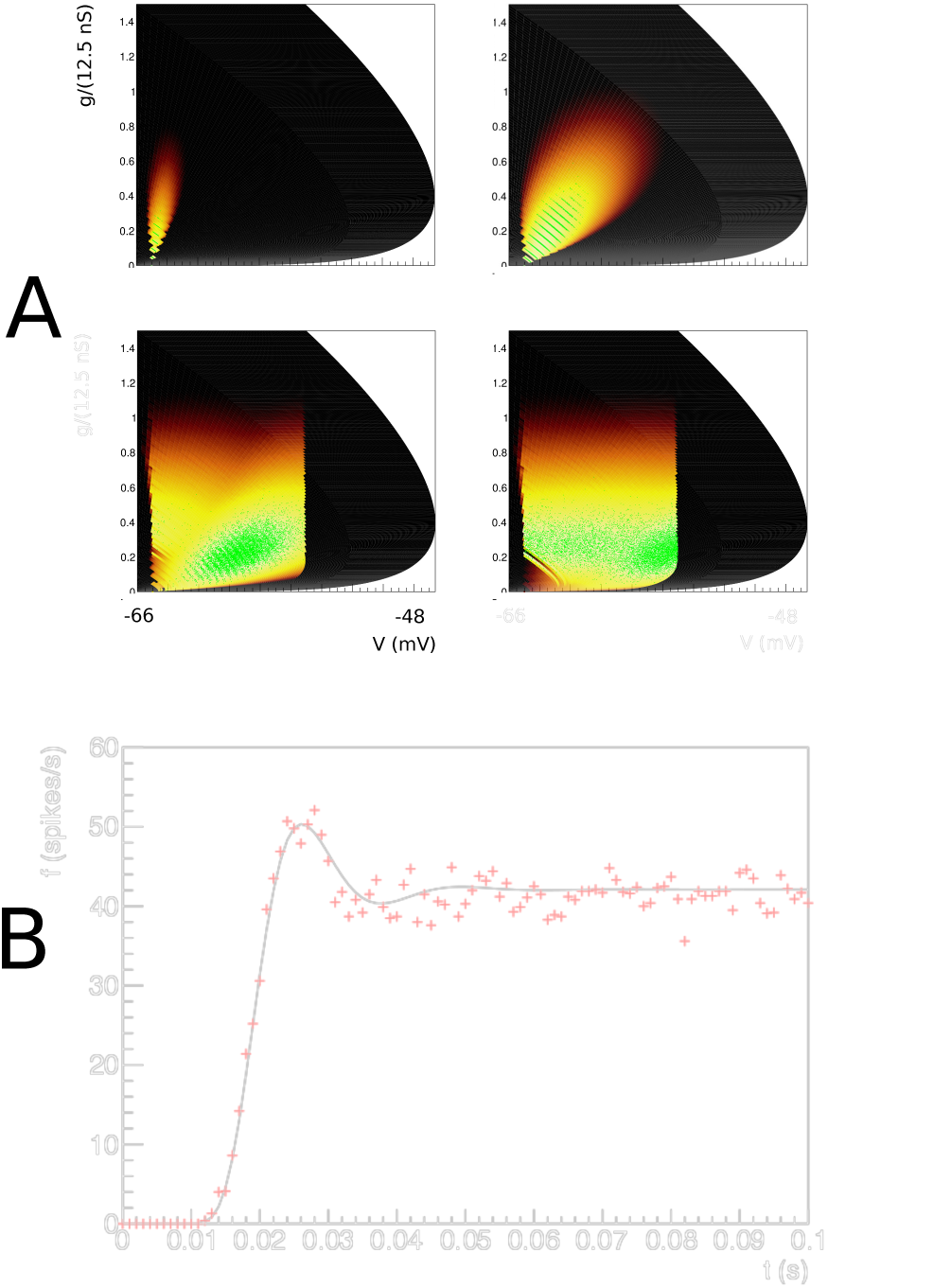
A: The evolution of the joint probability density function at four different points in time (1, 5, 15, 28 ms) in *(V,g)* space (*V* membrane potential, *g* conductance) for synaptic efficacy *J* = 0.05, Poisson generated input spike train with rate *v* = 1000 spikes per second. B: The resulting population firing rate, calculated from the fraction of mass crossing threshold per unit time as a solid black line. Spiking neuron simulation results shown by red markers. Onset and resulting firing rates are in agreement throughout. Unlike one dimensional neural models, conductance based models produce almost no overshoot.

The density is represented as a heat plot: the maximum density is white, lower density areas are shown as cooler colours from white through yellow to red. The color scale is logarithmic, so red areas represent very low probability. Figure 4 A) shows the evolution of the density of a population that was at equilibrium at *t* = 0 at four points in time *t* = 1,5,15 and 28 ms by which time steady state has been reached. We see probability mass moving mainly upwards under the influence of incoming spike trains. We will see that the mass ‘rotates’ in the direction of the threshold; and finally a steady state is realized: a state where the density profile has become stationary. We also have simulated a group of 10000 neurons and modeled incoming Poisson spike trains for each one. We keep track of their position in (*V, g*) space and represent their state at a given time as points in state space. The cloud of points clearly tracks the white areas of the density. The shot noise structure is clearly visible in the band structure early in the simulation where neurons are present at multiples of the synaptic efficacy, reflecting that some neurons have sustained multiple hits by incoming spike trains.

As neurons are moving through threshold, they themselves emit a spike and contribute to the response firing rate of the population, defined as the fraction of the population that spikes per time interval, divided by that time interval. We can therefore calculate the response firing rate from the amount of mass moving through threshold per unit time. We show the jump response of the population as a plot of populating firing rate as a function of time in Fig. 4 B. The firing rate calculated from the density matches that calculated from the Monte Carlo simulation very well. Interestingly, there is almost no overshoot in the firing rate, as also noted by Richardson (2004), who studied this system using Fokker-Planck equations. Although we study shot noise, in the absence of a fundamental scale in the *g* direction, the central limit theorem ensures that the marginal distribution in *g* is Gaussian within a few milliseconds. It is clear that the population disperse in the *g* direction and drifts towards the threshold relatively slowly. The absence of a barrier allows the dispersal of the population before it hits threshold, greatly reducing any overshoot in the firing rate, which is quite unlike one dimensional neural models, as we shall see in Sec. **Results: One Dimension**.

Let us contrast this with a simulation where we introduce a maximum conductance *g*_*max*_ = 0.8, which for simplicity we assume to be voltage independent. This then introduces a reflecting boundary at *g = g*_*max*_, and therefore introduces a scale by which an efficacy can judged to be small or large. As expected, probability mass is squashed against this boundary (Fig. 5 A) and has nowhere to go but laterally, in the direction of the threshold. Interestingly, the mass has not dispersed and clear groupings of mass huddled against the boundary can be observed. The traversal of the threshold by these groupings produces clear oscillations in the firing rate: a “ringing” effect. The firing rate jump response reflects the effect of the presence of a maximum conductance in state space.

**Fig. 5.**
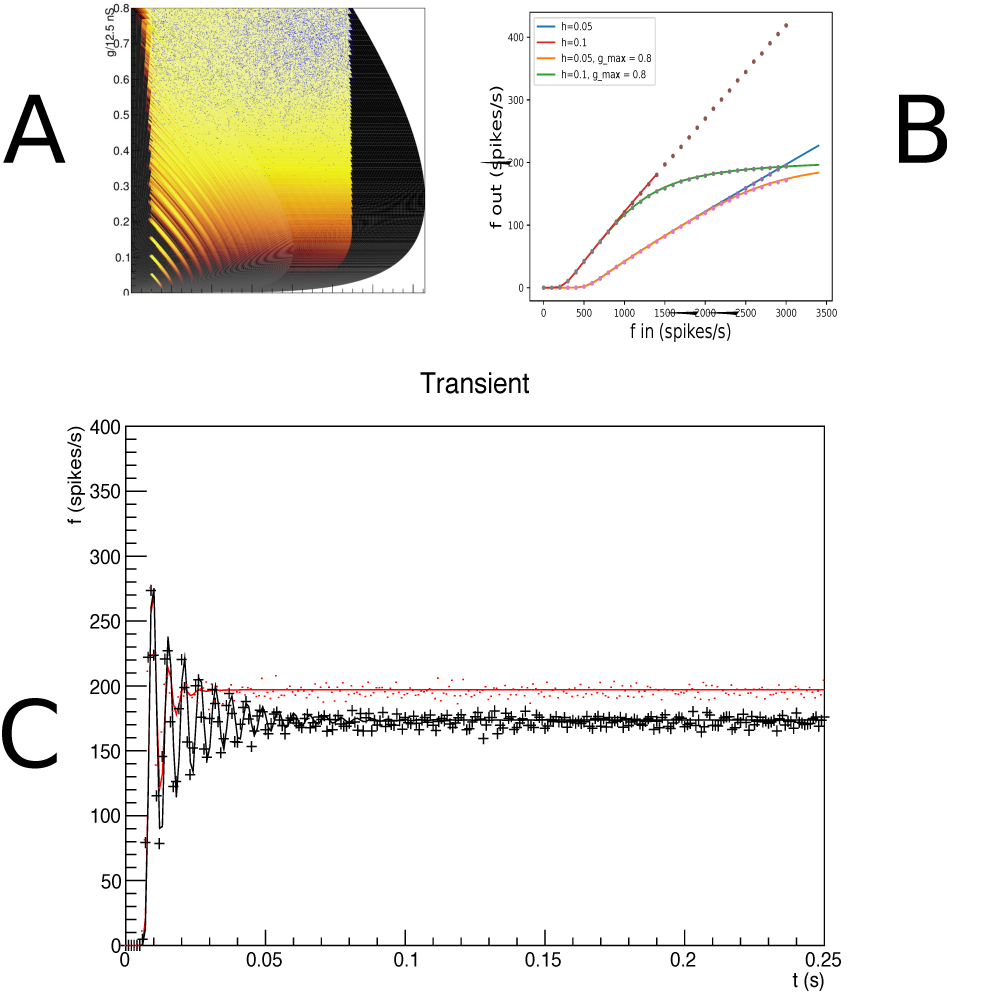
A: The density at *t* = 15 ms. Events are reflected against a reflecting conductance boundary. B: Gain curves for different input rates and synaptic efficacies. The maximum conductance clearly affects the shape of the gain curves, although at input rate 3 kHz for *J* = 1 mV the effect is moderate. C: The transient looks very different in the case where a maximum conductance is present: the “ringing effect” is much stronger. The cause can be seen in A: neurons have not had time to disperse before they are forced across threshold; clear groupings can be seen at the maximum conductance.

We run two simulations: one with and one without maximum conductance, but otherwise identical, and repeat this experiment for two different synaptic efficacies: *J* = 1 and 3 mV. Both simulations use an input rate of 3 kHz. In the case of no maximum conductance, probability mass can disperse in the *g* direction and mostly does so before arriving at the threshold. In Fig. 5 one sees that the introduction of a maximum conductance leads to a reduced response firing rate for high inputs. This can be interpreted as the population unable to respond to an increase of input once the majority of its ion channels are already open. Fig. 5 shows that the firing rates of Monte Carlo simulations and our method agree over the entire range of input.

Even when the effects on the response firing rate are moderate, the transient dynamics can be radically different. For an efficacy *J =* 1 mV and and input rate *v*_*in*_ = 3 KHz, the firing rates for maximum conductance, compared to no maximum come out as 175 Hz vs 195 Hz. In Fig. 5 C we show the response firing rate as a function of time. The result for the unrestrained conductance is given by the red line, which despite the high output firing rate still almost produces no overshoot. When we restrict the maximum conductance we see a somewhat reduced firing rate but a pronounced transient response (“ringing”) which persists much longer than for an unrestrained conductance. It is striking to see that the reintroduction of a barrier in state space results in pronounced transients. In both cases, the calculated firing rates agree well with Monte Carlo simulation. We attribute this ringing to a geometrical effect: the introduction of a barrier in the direction of where the stochastic process is pushing neurons.

### One Dimension: Leaky-and Quadratic-Integrate-and-Fire Neurons and Size Effects on the Transition Matrix

Although these model neurons are characterized by a single dimension - the membrane potential - they can be viewed as a two dimensional model that is realized in a single strip, and where transitions take place between one bin in potential space to another. This is completely equivalent - in implementation and concept - to the geometric binning method introduced independently by de Kamps [12] and Iyer et al. [11], with one exception: the generation of transition matrices by Monte Carlo. In one dimension it is not necessary to use Monte Carlo generation: the transition matrix elements can be calculated to an arbitrary precision because in one dimension the geometrical problem outlined in Sec. **Materials and Methods: Handling Synaptic Input** is much simpler and can be solved by linear search. It is clear that unlike the 2D case, it is straightforward to find the exact areas covered by translated bins, and hence no Monte Carlo generation process is required.

Nevertheless, it is interesting to use this method. The transition matrix generation for the 2D case is relatively expensive, and as precision scales with the square root of the number of events it is interesting to see how few we can use in practice without distorting our results. The answer is: surprisingly few. As benchmark we set up a population of LIF neurons with membrane constant *τ* = 50 ms, following [5], and assume that each neuron receives Poisson distributed spike trains with a rate *v* = 800 Hz. We assume delta synapses, i.e. an instantaneous jump in the postsynaptic potential by a magnitude *h =* 0.03, with the membrane potential *V ∊* [−1,1), i.e. we use a rescaled threshold potential *V* = 1. The grid is generated with a time step Δ*t* = 0.1 ms, and is shown in Fig. 6 B.

**Fig. 6.**
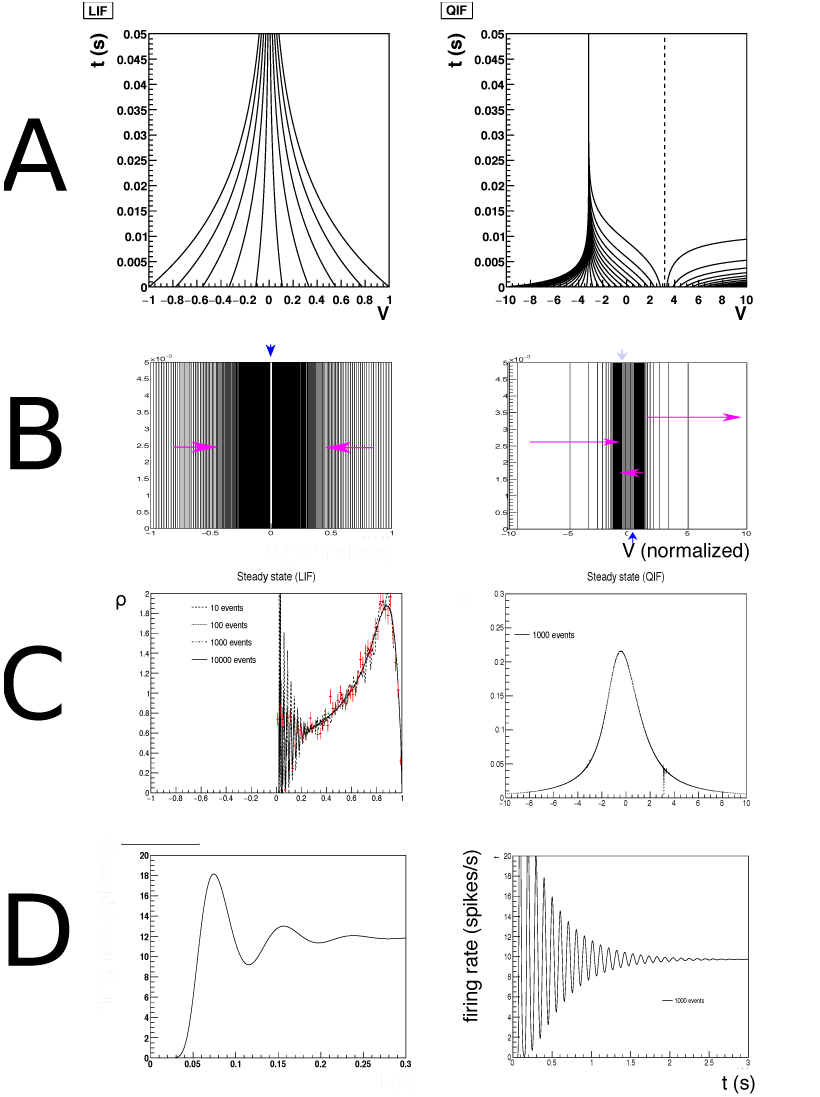
A: Characteristics for leaky-(left) and quadratic-integrate-and-fire (right) neurons. B: The resulting grids for both neuron types. C: Typical steady state densities, strongly influenced by the shape of the grid, and ultimately the neural dynamics. D: a typical jump response of the firing rate. For comparable output frequencies, QIF neurons “ring” for longer, which we attribute to a closer grouping of probability mass.

The simulation results are shown in Fig. 7 and replicates earlier work [5,12]. The use of a finite number of points in the Monte Carlo process used for the generation of transition matrices generates random fluctuations with respect to the true values. The effect of these fluctuations is clearly visible in the shape of the density profile, and only for *N_point_ =* 10000 the profile is as smooth as in earlier results where we calculated the transition matrix analytically. How bad is this? To put these fluctuations into perspective, we used a direct simulation of 10000 spiking neurons and histogrammed their membrane potential at a simulation time well after *t =* 0.3 s, so that they can be assumed to sample the steady state distribution. In the figure, they have been indicated by red markers. Comparing the results we see that the fluctuations for *N*_*point*_ = 10 are comparable to those of a Monte Carlo simulation using a sizeable population of 10000 neurons. Moreover, in the population firing rates the finite size effects are almost invisible. This is somewhat surprising, but a consideration of the underlying process that generates the firing rate explains this. Neurons are introduced at equilibrium and will undergo several jumps before they reach threshold. The finite size effects of the Monte Carlo process induce variations in those jumps in different regions of state space, but these fluctuations are unbiased and will average out over a number of jumps. So neurons will experience variability in the time they reach threshold, but this variability does not come in the main from fluctuations in the transition matrix elements. It should be emphasized that the transition matrices are a quenched source of randomness, because transition matrices are fixed before the simulation starts. So although ultimately caused by finite size effects, their contribution is different compared to the unquenched finite size effects that can be seen in the population of 10000 neurons.

**Fig. 7.**
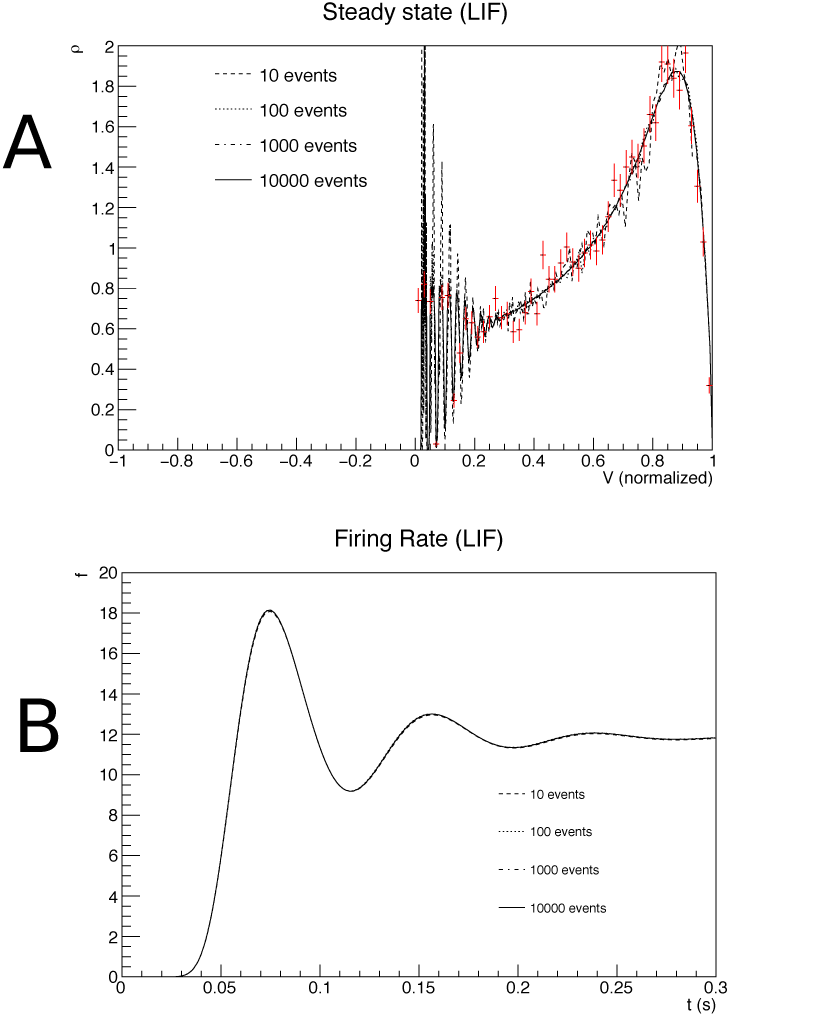
A: The steady state density as a function of membrane potential. B: The firing rate as a function of time, for transition matrices that were generated for different values of *N*_*point*_, the number of events used in the Monte Carlo generation of transition matrices.

It is instructive to look at some examples because it highlights strengths and weaknesses of the method in terms of familiar results. In Fig. 6 A, the characteristics of both neural models are given. In Fig. 6 B the state space of LIF (left) and QIF neurons (right) are shown, at lower resolution than used in simulation to elucidate the dynamics. Rather than with numbers which would be unreadable at this scale, we indicate the direction in which cell numbers increase, and therefore the direction in which neural mass will move, by arrows. One can see that the LIF neuron is comprised of two strips, and the QIF neuron of three, where the arrows indicate in which direction the cell numbers are increasing. In the LIF grid, there is one stationary bin, in the QIF there are two. They are represented as separate stationary cells, covering the space between the strips, indicated by the blue downward pointing arrows.

In Fig. 6 C we consider the steady state of LIF (left) and QIF neurons (right) after being subjected to a jump response of Poisson distributed spike trains starting at *t* = 0 (LIF: *v*_*in*_ = 800 spikes/s *J* = 0.03 (normalized w.r.t. threshold; QIF: *J* = 0.05)). The shape of the characteristics and therefore of the grid clearly reflect their influence on the steady state density distribution. The output firing rate (Fig. 6 D) shows the clear “ringing” in the transient firing rate that is mostly absent in conductance based neurons. Again, this can be interpreted geometrically: the stochastic process pushes neurons in the direction of a threshold, but they reach it without having had the opportunity to disperse. Decorrelation only happens after most neurons have gone through threshold at least once. It is also interesting to see that for comparable firing rates the ringing is much stronger for QIF than for LIF neurons. We also interpret this as a geometrical effect: the effective threshold for QIF neurons is *V* = 3 (normalized units), not 10, as neurons with a membrane potential above 3 will spike. It is clear from Fig. 6 D that compared to LIF neurons, QIF neuron bulk up close to the threshold and are constrained more than their LIF counterparts, thereby making it harder to decorrelate before passing threshold.

For reference, in Fig. 8 we show that the method accurately reproduces results from the diffusion limit, as well as generalizes correctly beyond it. If one uses a single Poisson spike train to emulate a Gaussian white noise input, employing the relationship:

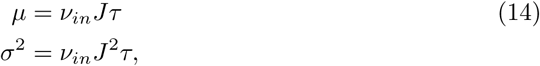

one can use our method to predict the steady state firing rates as a function of *J*, the synaptic efficacy and *v*_*in*_ the rate of the Poisson process for given membrane constant *τ*. Organizing the results in terms of *μ* and *σ*, as given by Eq. 14, one expects a close correspondence for low *σ*, since Eq. 14 leads to small values of *J* compared to threshold. One expects deviations at high *σ*, where *J* does not come out small. Fig. 8 shows that this is indeed the case when firing rates are compared to analytic results obtained in the diffusion approximation. Our method produces the correct deviations from the diffusion approximation results, and agrees with Monte Carlo simulation. Elsewhere [12], we have shown that diffusion results can be accurately modeled using two Poisson rates for high *σ*.

**Fig. 8.**
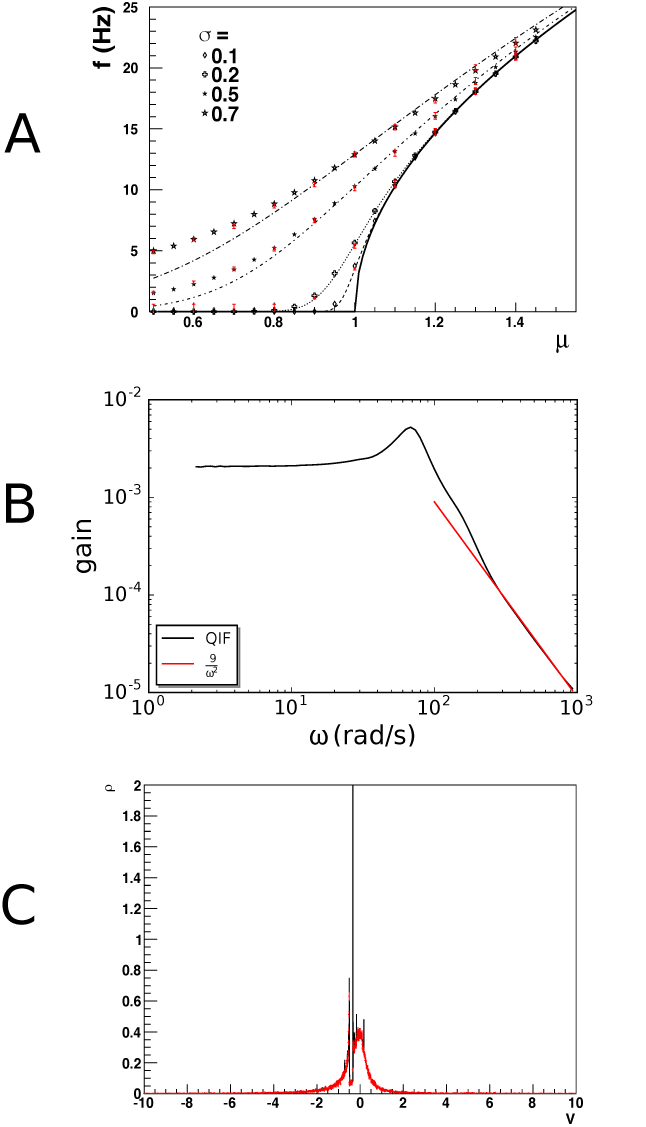
A: Gain curve for quadratic-integrate-and-fire neurons. Population density techniques handle deviations from the diffusion approximation correctly: when one tries to emulate Gaussian white noise with a single Poisson spike train input, deviations are expected at high values of *σ* as synaptic efficacies are forced to be large. The dashed lines give the diffusion approximation, black markers the prediction by our method and red bars Monte Carlo results, which agree with each other, but deviate from the diffusion prediction. B: The frequency spectrum shows the expected 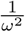 dependency for high frequencies. C: the delta peak of a coherently firing group of neurons in correctly represented; the decrease in partial synchrony of the population is modeled correctly over long times.

In Fig. 8 B we replicate the gain spectrum for QIF neurons and show that the high frequency dependence falls off as 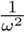 as predicted by Fourcaud-Trocme *et al.* [24]. These results reaffirm that our method accurately predicts results within and beyond the diffusion limit, and that a substantial body of existing literature can be seen to be a special case of our method.

Figure 8 C shows a population of QIF neurons that fire in synchrony at *t* = 0, undergoing a slow decorrelation by low rate Poisson input spike trains. The neurons have all been prepared in the same state, and therefore are at the same position in state space. We use *F(V)* = *V*^2^ + 1, so these neurons are bursting, as the current parameter is larger than 0, and there are no fixed points. Neurons that receive an input spike leave the peak and travel on their own through state space. This results in a very complex density profile, where the initial density peak is still visible after 1s. Such a peak would have diffused away rapidly in a diffusion limit approximation. Monte Carlo events in red markers show that the density profile is not a numerical artefact, but reflects the complexity of the density profile.

### Adaptive-Exponential-Integrate-and-Fire Neurons

We consider the AdExp model as presented by Brette and Gerstner [25], which describes individual neurons by the following equations:

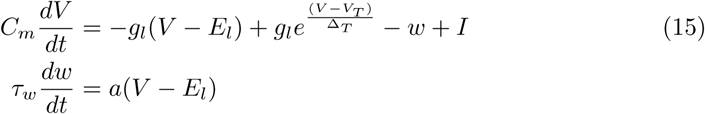

Upon spiking, the neuron is reset to potential *V*_*reset*_ and increases its adaptivity value: *w* → *w* + *b*. Here *C*_*m*_ is the membrane capacity and *g*_*l*_ the passive conductance. *V*_*T*_> is the value at which a neuron starts to spike; the spike dynamics is controlled by Δ_*T*_. The numerical values of the parameters are summarized in Table 2 and are taken from [25].

An overview of the state space is given in Fig. 9 A. At *w* = 0 the dynamics is as expected, a drive towards the equilibrium potential that suddenly reverses into a spike onset at higher values of *V*, essentially producing an exponential-integrate-and-fire neuron. At high *w* two effects conspire to make the neuron less excitable: the equilibrium potential is lower and the drive towards this equilibrium is stronger for a given value of *V.* At low *w* values, the opposite happens: the equilibrium value is higher, closer to threshold, and below equilibrium there is a stronger depolarizing trend making the neuron more excitable. Interestingly, at hyperpolarization the system does not only respond by driving the membrane potential back towards equilibrium potential, but also downwards.

**Table 2.** Parameters for the AdExp model as given in [25]

| Quantity | Value |
| --- | --- |
| $C_m$ | 281 pF |
| $g_l$ | 30 nS |
| $E_l$ | -70.6 mV |
| $V_T$ | -50.4 mV |
| $\Delta_T$ | 2 mV |
| $\tau_w$ | 144 ms |
| $a$ | 4 nS |
| $b$ | 0.0805 nA |

**Fig. 9.**
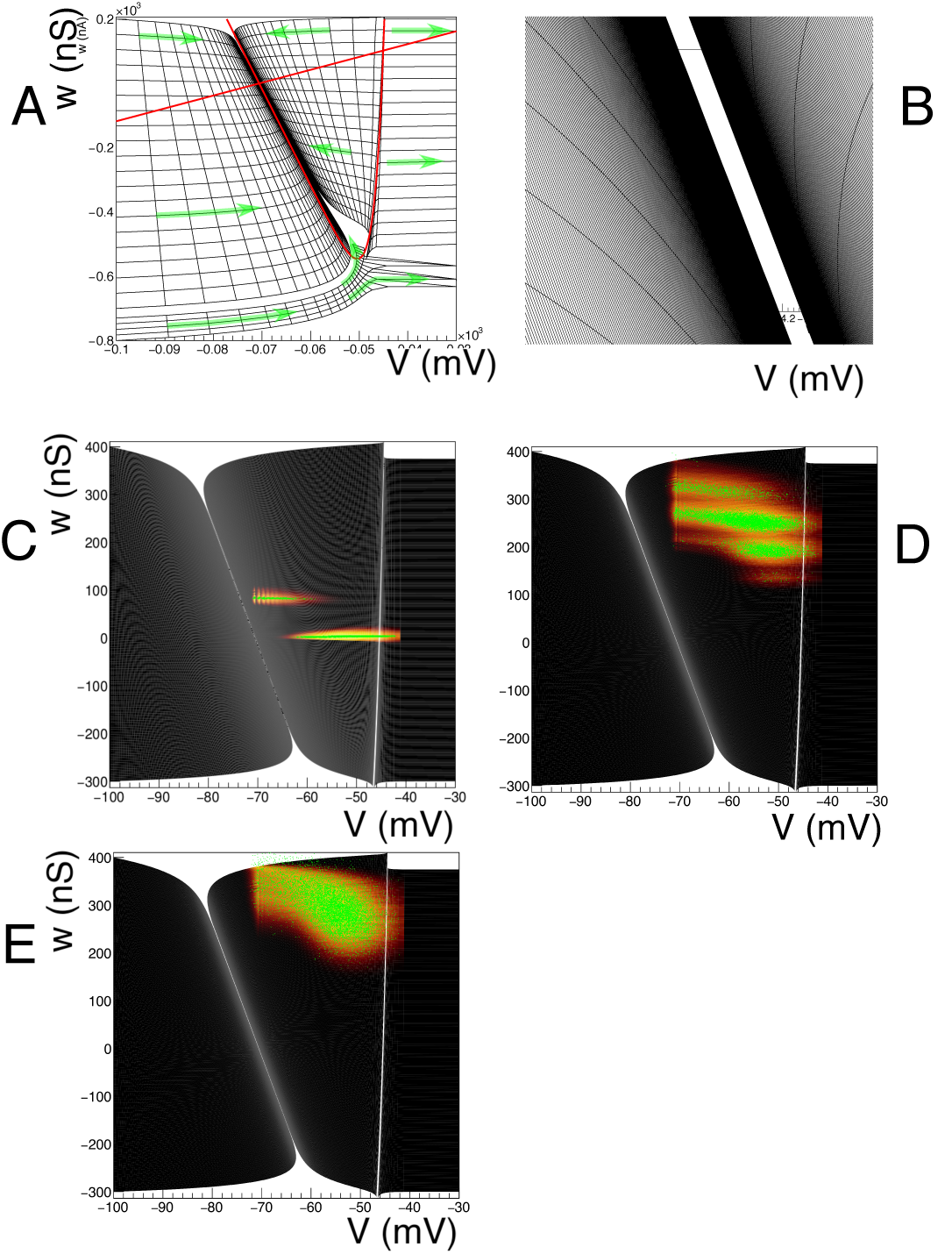
A: Overview of state space for the adaptive-exponential-integrate-and-fire neuron. B: a detail of a realistic mesh near the equilibrium potential. C-E: evolution of the probability density at *t* = 0.01, 0.1, 0.4s. The input (switched on at *t* = 0) is a Poisson distributed spike train of 3000 spikes/s, delta synapses with efficacy *J* = 1 mV.

There are two critical points, the equilibrium point (*E*_*l*_,0) and a saddle point in the top right. They are at the crossing of two nullclines: the w-nullcline is a straight line, whereas the V-nullcline follows a strongly curved trajectory, which is close to the stable manifold of the saddle point in a substantial part of state space. Below (to the right) the stable manifold neurons spike, regardless of where they are initially, while above (to the left) of the stable manifold neurons converge to the equilibrium, but how, and how long this takes is strongly dependent on the initial conditions. This model is the first to require a judicial treatment of the grid boundaries.

Let us examine the the equilibrium point first. The exponential build-up of cells observed in one dimensional models occurs here as well, but here it is not a good idea to introduce a fiducial cut and cover the remaining part of state space with a cell. The inset of Fig. 9 B shows that equilibrium is reached much faster in the *V* direction, than in the *w* direction. This is a direct consequence of the adaptation time constant *τ*_*w*_ being an order of magnitude larger than the membrane time constant *τ* ≡ *C*_*m*_/g_*l*_. For high *w*, mass will move downwards along the diagonal, until low values of *w* are reached, as is demonstrated by the left inset of Fig. 9. A long, but very narrow region separates different parts of the grid. What to do? First, we observe that the offending region is essentially forbidden for neurons: for most neurons starting from a random position in state space it would take a long time (of the order of 100 ms) to approach this no man’s land. At the input firing rates we will be considering, neurons will experience an input spike well before running off the strip, so essentially only noise can place neurons there. If we forbid this, by allocating events that are translated into the cleft between the two grid parts to the cells in the grid that are closest to it along the projection of the jump, we guarantee that no probability mass will leak out of the grid. Mass that reaches the end of the strips will be placed in a reversal bin, like the one dimensional case. Mass on the left of the side of the cleft will move in the same direction as that on the right side of the cleft. By using Euclidean distance projected along the jump direction, we minimize the bias due to this procedure, although we may artificially introduce a small extra source of variability.

On the right hand side, the stable manifold almost coincides with the *V* nullcline, resulting in a very narrow region of dynamics in the vertical direction. Immediately outside neurons rapidly move away laterally. This part of the grid is created by reversing the time direction, integrating towards the stable manifold. The grid strongly deforms here: cell area decreases rapidly and even small numerical inaccuracies will lead to cells that are degenerate. We use cell area as a stopping criterion. The last cells before breaking off are extremely elongated. The spike region is also created by reversing the time direction. Again, we conclude that the cleft is a forbidden area: a small fluctuation in the state variable will cause a neuron to move away rapidly. Our main concern, again, is neurons that are placed into this cleft by the noise process. Again, we move neurons to the closest cell next to the cleft in the jump direction. This is reasonable, since natural fluctuations would put them there soon anyway. Effectively we have broadened the separatrix a little bit, but we still capture the upwards (for high *w* - past the saddle point: downwards) movement close to the stable manifold.

In Fig. 9 C-E the evolution of a population in (V, *w)* space is shown at three different points in time: *t* = 0.05, 0.1 and 0.4 s. Figure 9 C shows the input spikes pushing the state towards threshold, and a small number of neurons have spiked. They re-emerge at the reset potential, but with much higher *w*, due to spike adaptation. This is determined by the b parameter of the AdExp model. Close to the reset potential the banded shot noise structure, due to the use of a delta-peaked synaptic efficacy, is visible. The steady state is reached after approximately 400 ms. The population stabilizes at high w values, and the bulk of the population is clearly well below threshold, due to stronger leak behavior at these values of *w*. In sub figure E there is a minute deformation of the density, due to the limits of the grid, and density heaps up here, but the fraction of probability mass affected is negligible. Monte Carlo events, indicated by the dots, are not restricted to the grid and some fall outside the grid.

The firing rate response corresponding to the population experiencing an excitatory input (Fig. 9 C-E) is given in Fig. 10 A. Again, agreement with Monte Carlo simulation is excellent, we are able to study the relative contributions of current-and spike-based adaptation to the firing rate. We can easily simulate neurons with current-but not spike-based adaptation by not incorporating the jump in *w* after reset; while ignoring all forms of adaptation can be done by simply using a 1D grid and ignoring values of *w* ≠ 0. The vast difference between adaptive neurons and non-adaptive neurons is also reflected in the gain spectrum. Figure 10 B shows the gain spectrum of a (non-adaptive) exponential-integrate-and-fire neuron and a neuron that has a constant rate of adaptation due to the background rate upon which the small sinusoidal modulation has been imposed. The difference between the adaptive and non-adaptive neuron is considerable. Both neurons show 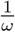 dependence in the high frequency limit, as is expected for exponential neurons [24]. (Fig. 10 A shows that the shape of the spike, which is reflected in the large cells on the right of the grid is independent of *w*.)> It is clear that a meaningful time-independent gain function cannot be chosen, so that it is not possible to develop linear response theory.

**Fig. 10.**
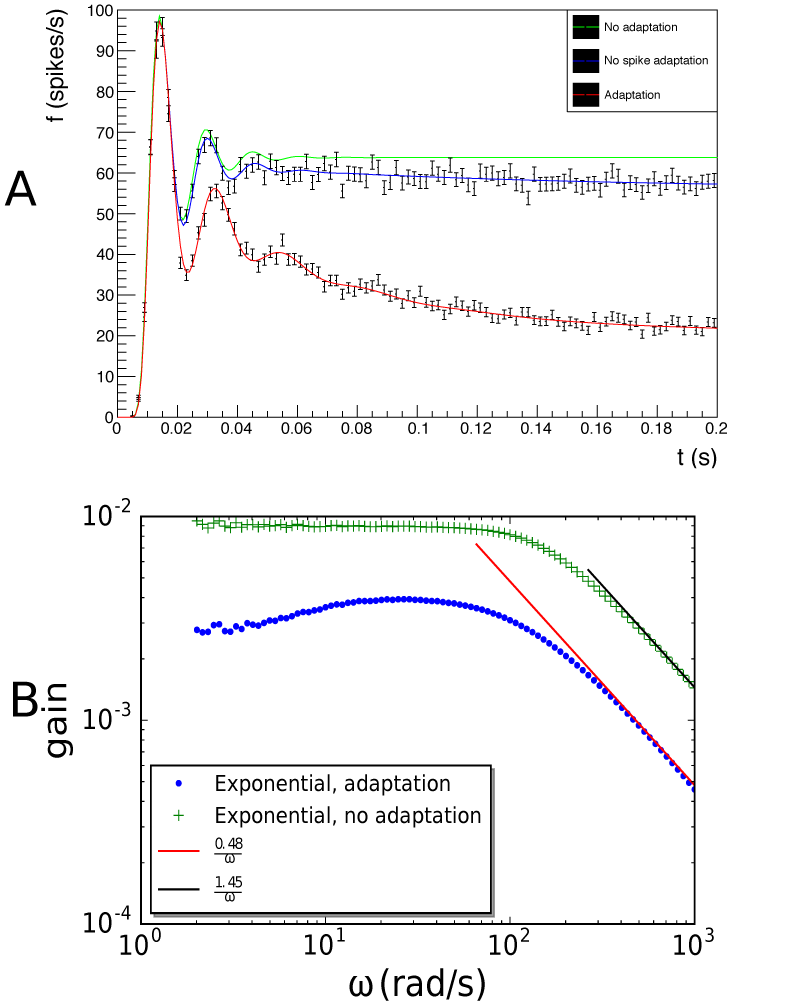
A: compares the response for three cases: no adaptation; only current adaptation; and both current-as well as spike-based adaptation is included. B: The gain for a small sinusoidal input modulated on a background input as function of frequency, for no adaptation and AdExp with current-and spike-based adaptation. Both spectra show the 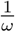 dependency expected of an exponential-integrate-and-fire neuron, as the spike shape, represented by the grid at high *V* values is independent of *w*. However, the numerical difference between the cases is vast.

It is interesting to observe the marginal distributions - in Fig. 11 we show the marginal distributions, together with the joint distribution. The distribution in *V* looks remarkably like that of an LIF neuron, except near the threshold, where the spike region, which is not present for LIF, flattens the density. The *w* distribution suggests a much stronger overlap than the joint distribution, which shows a clear separation. It is clear that, had the three density blobs been oriented more diagonally, the marginal *w* distribution would have shown a single cluster.

**Fig. 11.**
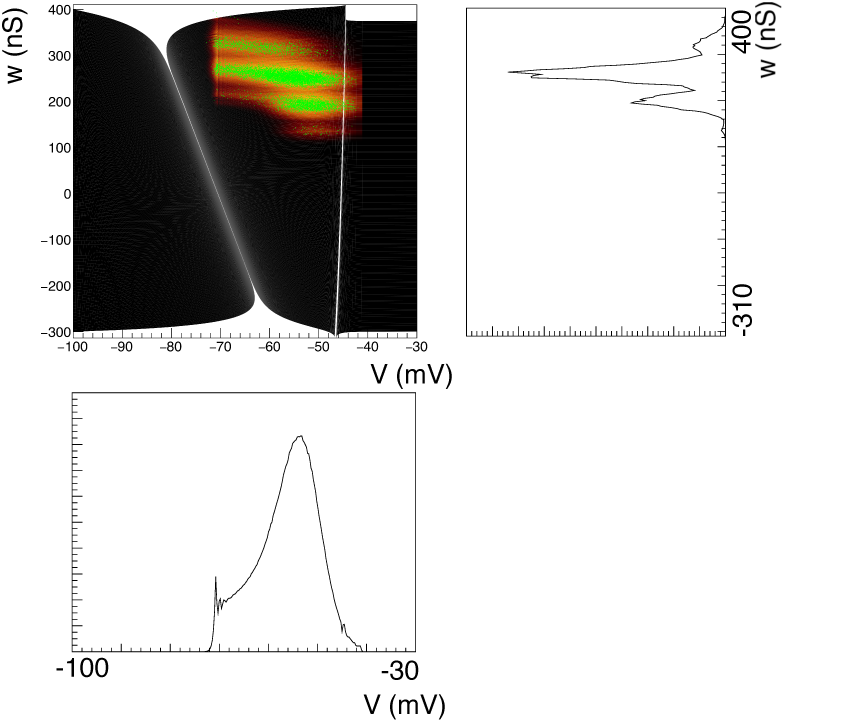
Comparison of the joint distribution function of the AdExp neuron with the marginals in *V* and *w*. The marginal *w* distribution still reveals four clusters, but the joint distribution reveals them as being far better resolved than one would judge from the marginals.

### Frequency-dependent Short-term Synaptic Dynamics

Vasilaki and Giugliano have studied the formation of network motifs [22], using both microscopic spiking neural simulations and mean-field approximation. In their mean-field simulations they considered both spike-timing dependent long-term plasticity, and frequency-dependent short-term dynamics, where they use a version of the Tsodyks-Markram synapse [26]. The short-term dynamics is of interest because it introduces something we have not considered before: the magnitude of the jump being dependent on the position of where the jump originates. Following [22], if *G*_*ij*_ defines the amplitude of the postsynaptic contribution from presynaptic neuron *j* to postsynaptic neuron *i*, then this is considered to be proportional to the amount of resources used for neurontransmission *u*_*ij*_*r*_*ij*_ and to their maximal availability *A*_*ij*_, so

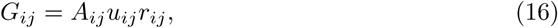

where *r* relates to the recovery and *u* to the facilitation of synapses, and the time constants *τ*_*rec*_ and *τ*_*facil*_ are different for facilitating and depressing synapses. They describe frequency-dependent short-term synaptic dynamics by:

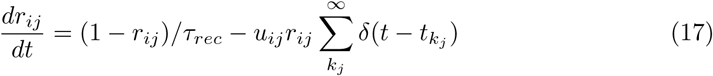

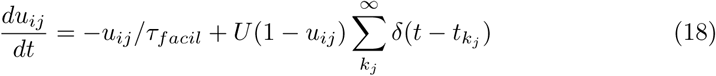

From now on, we will drop the indices *ij* and just refer to a single connection. In the simulation below we will use *τ*_*rec*_ = 0.1 s and *τ*_*facil*_ = 0.9 s and study a population of facilitating synapses (Vasilaki and Giugliano used *τ*_*rec*_ = 0.9s, *τ*_*facil*_ = 0.1 s for depressing synapses.) *U* is a fixed constant, for facilitating (depressing) synapses *U* = 0.1(0.8). Equation 17 expresses that an individual synapse is subject to deterministic dynamics, and that upon the arrival of a spike at time *t*_*k*_ both *u* and *r* undergo a finite jump, whose magnitude is dependent on the current value of *u* and *r*. Equation 2 describes this situation, when the following transition probabilities are introduced:

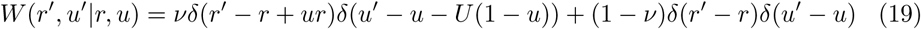

We have to modify the process of generating our transition matrices: now for each quadrilateral cell (*p*, *q*), we determine the centroid (*u*_(*p,q*)_, *r*_(*p,q*)_) and we determine the covering set by defining

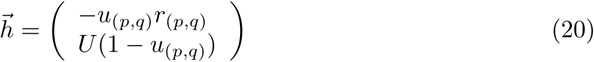

and determining the cover set as before. The jump now becomes cell dependent.

It is easy to cover almost the entire state space. In Fig. 12 A we show the grid. In Fig. 12 B, we show the sample path of three synapses, assuming that the presynaptic firing rate *v* = 5 Hz. In C-F we show the evolution of a population of synapses. The influence of the step size which increases in the *r* (horizontal) direction with *u* and *r*, but decreases in the *u* (vertical) direction with *u*. There is good agreement with Monte Carlo simulation throughout.

With the joint distribution available, it is possible to use Eq. 16 and calculate the distribution of *G*_*ij*_ or its expectation value.

### Fitzhugh-Nagumo Neurons

We consider the well-known Fitzhugh-Nagumo neuron model [27], which is given by:

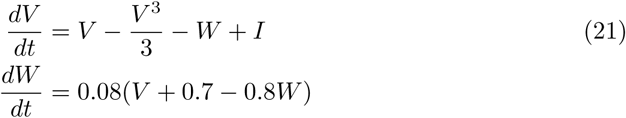

It is an attractive neuron model as it captures many properties of the biologically realistic Hodgkin-Huxley neuron, while being much more tractable - being reduced to two dimensions aids greatly in analysis and visualization. The two variables are a nondimensionalized voltage-like variable *V* and a recovery variable *W*. Also we note a variable *I* representing a constant external current.

When *I* = 0, there is a stable equilibrium point at ≈ (- 1.199, −0.624) corresponding to a resting state. As *I* increases, the system undergoes a Hopf bifurcation to a stable limit cycle around an unstable equilibrium. (Increasing *I* further leads to a stable fixed point at positive *V* and *W* termed “excitation block”.) In this paper, we will consider an intermediate value *I* = 0.5 in order to demonstrate how our method can be used on systems with limit cycles.

**Fig 12.**
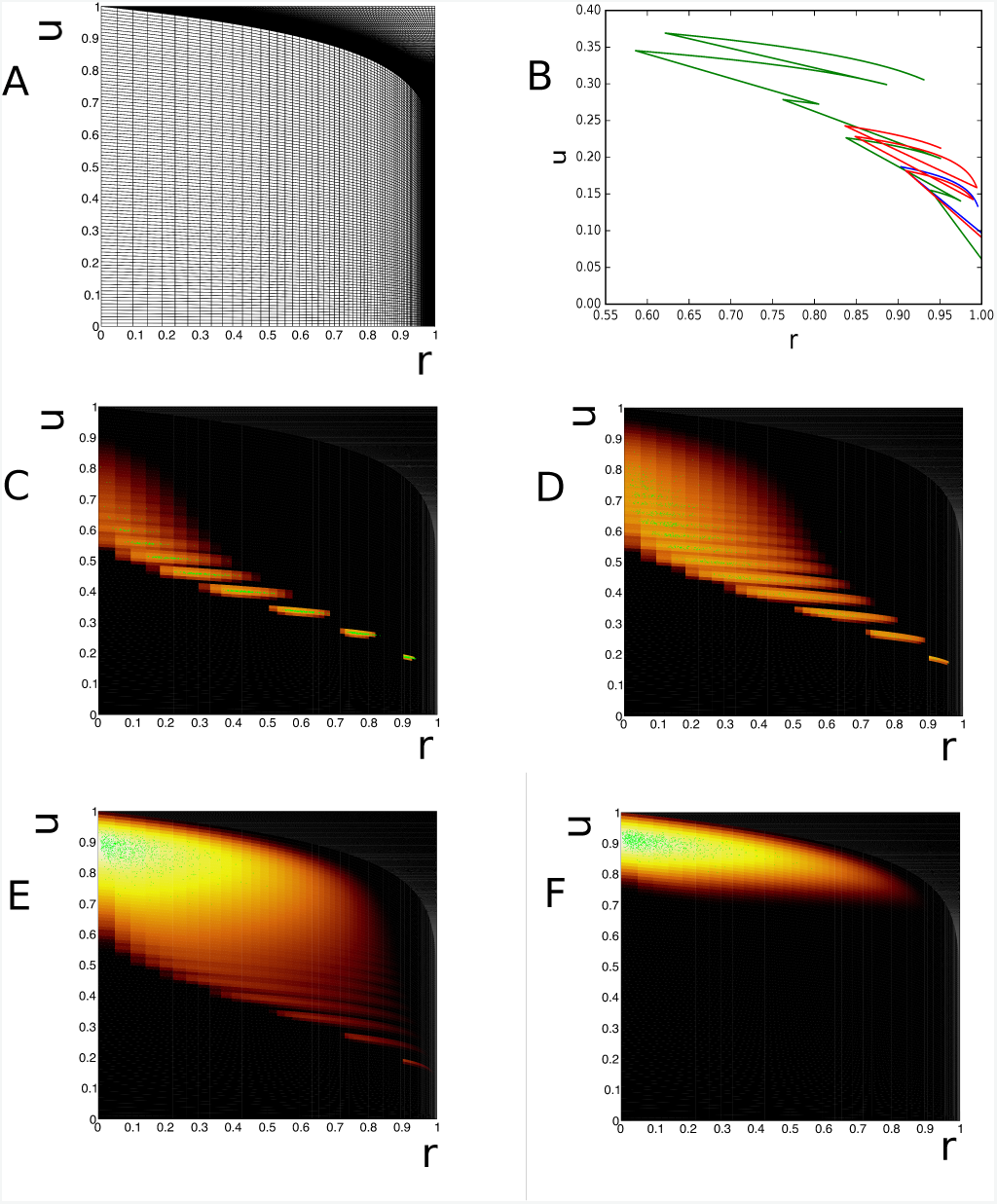
Evolution of the state of a population of facilitating Tsodyks-Markram synapses. A: Grid. B: Sample path of individual synapses. The state dependency of the jump size is clearly visible. C-F: evolution of density over time.

We simulate white noise by providing the system with both inhibitory and excitatory noisy input with a high rate and low synaptic efficacy, and successfully capture the diffusion of probability in a neighbourhood around the limit cycle (Fig. 13 A-D for *t* = 5, 10, 50 and 1000 s, *J* = ±0.02, *v* = 20 spikes/s). It is interesting to study a purely excitatory input with large synapses (*J* = 0.1, *v* = 2 Hz). This leads to a deformed limit cycle, shifted towards higher *V*. This is expected as the net input now is *I* = 0.7. The band is also broader, as one would expected as higher values of synaptic efficacy imply larger variability.

Another case we consider is noisy inhibitory input (Fig. 13 F). As we would expect, the system is effectively driven back below the bifurcation to a stable equilibrium, although we still see some variance-driven probability follow a limit cycle that differs considerable from the original limit cycle. We can understand this by converting the noisy input into zero-mean noise and a steady inhibitory current, and looking at the streamlines of the system with these parameters instead. As seen in Fig. 14 D, while all the the trajectories converge to the fixed point, those starting on the right side of phase space first increase *w* until they reach the right branch of the cubic nullcline, then follow a path close to the limit cycle to return to the fixed point. It is interesting to see that the method captures limit cycles that do not coincide with the limit cycle of the original grid.

**Fig 13.**
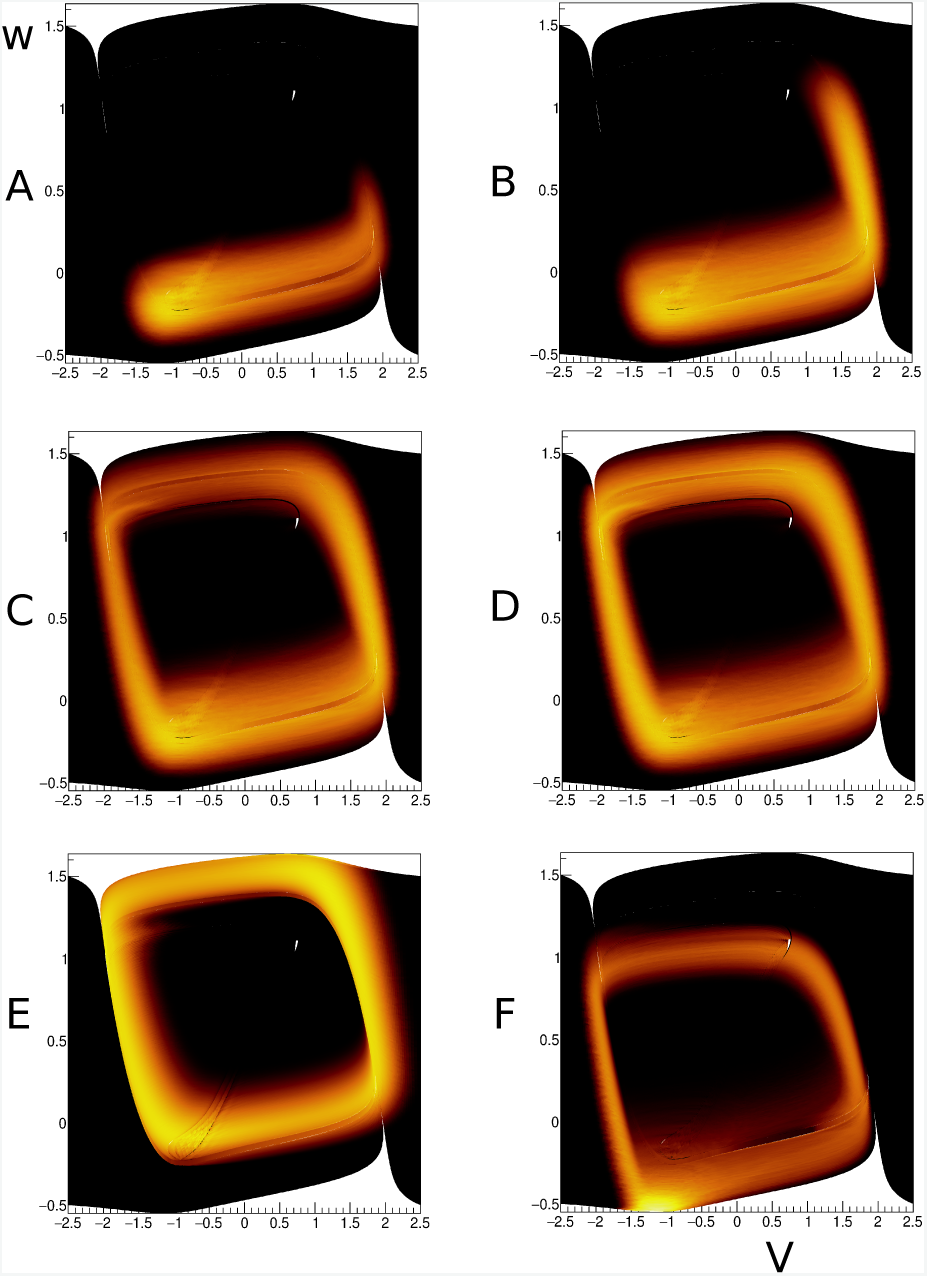
Box initially at rest on sled sliding across ice.

A-D: Evolution of the density at *t* = 5,10, 50 s an steady state for a diffusive input (*J* = ±0.02, *v* = 20 spikes/s). E: excitatory input for a weak rate, but with large synaptic efficacy (J = 0.1, *v* = 2 spikes/s). F: inhibition captures most neurons at the fixed point, a weak ghost cycle of neurons that escape by fluctuation is visible, but is considerably displaced compared to the standard limit cycle.

Rabinovitch and Rogachevskii [28] describe the two “vertical” sections of the path to be transient attractors (T-attractors) separated by a diagonal transient repeller (T-repeller) (alternatively, a separatrix [29]) close to the central branch of the cubic nullcline. Trajectories close to each other but starting on different sides of the T-repeller separate rapidly before eventually reaching the same steady state, which creates considerable problems in creating the grid (see Fig. 14 E). The authors perform a detailed analysis of the system by extending the notion of isochrones from limit cycles to excitable systems. We note that their isochrones are similar in character to the lines in our grid perpendicular to the streamlines of the system.

Next, we outline some of the numerical subtleties involved in generating the computational grid for the Fitzhugh-Nagumo model. Following the procedure from Sec. **Materials and Methods**, one can attempt to generate a grid by starting with a set of initial conditions, and solving the differential equations of the system forwards in time to obtain a set of trajectories (or integral curves). Each pair of trajectories then has a strip between them and the individual cells are obtained by dividing the strip into equal-time bins. However, in a system with a limit cycle, if we start with initial conditions outside the limit cycle, we see that the trajectories generated from them converge onto the limit cycle. Moreover, it is impossible to obtain trajectories inside the limit cycle from outside the limit cycle, and vice versa. This means that we have to handle the limit cycle, outside, and inside, as separate sections of the plane.

**Fig 14.**
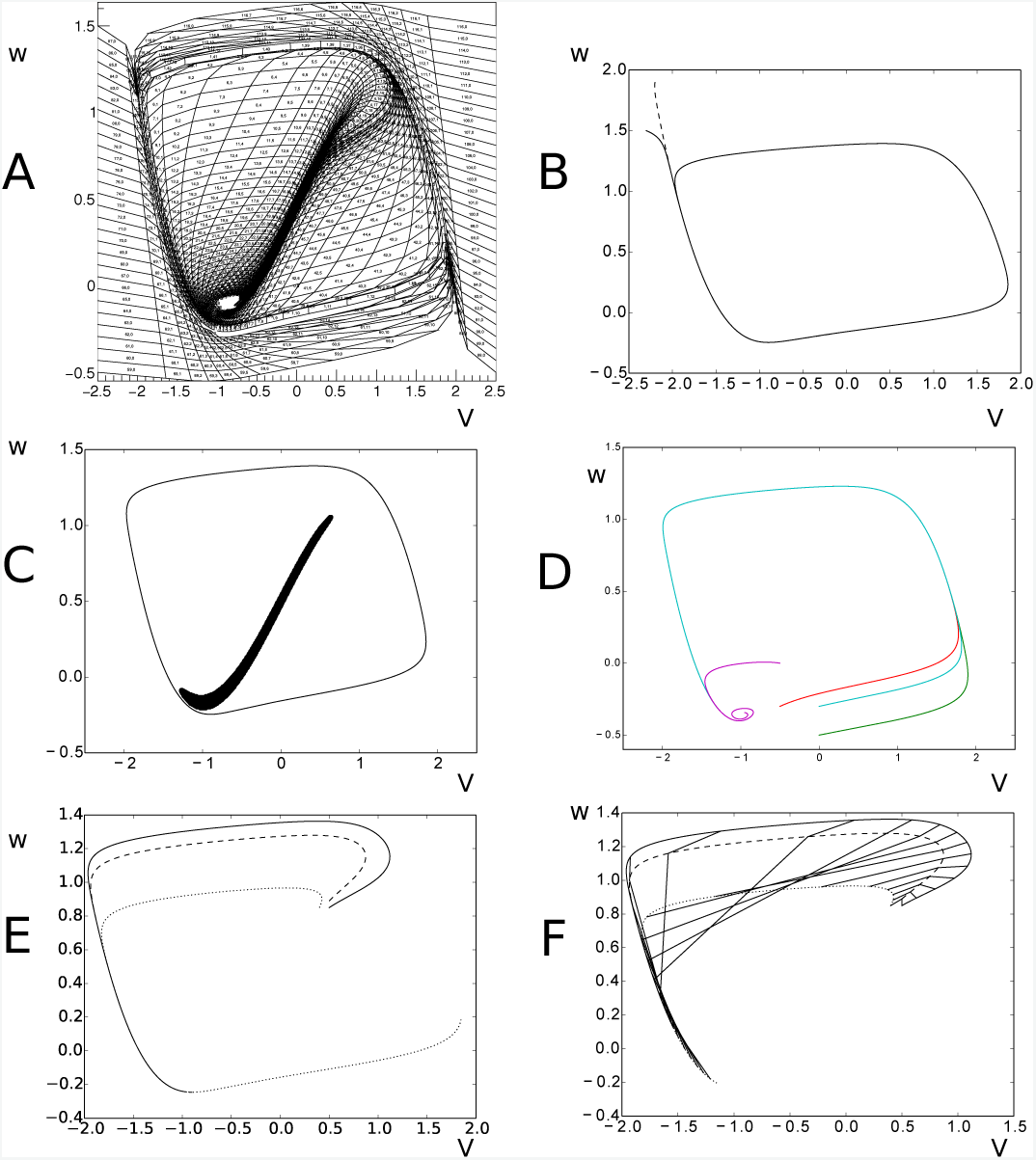
A: The numbered mesh provides a coarse overview of the dynamics of the system, whereas we run our simulations using the much finer mesh. B: Different trajectories in the Fitzhugh-Nagumo model (as shown by solid and dashed lines) can merge before reaching the limit cycle. C: The region of the Fitzhugh-Nagumo model tiled with stationary cells; limit cycle is shown for reference. D: Sample trajectories in the Fitzhugh-Nagumo model in the parameter regime where the system tends to a fixed point. E: Trajectories that begin close to each other have the potential to diverge rapidly. F: Attempting to build cells from these trajectories can lead to “stretched” cells that intersect other cells.

Since the limit cycle is a one-dimensional object with zero width, we have to artificially define a small width around it. We then choose sets of initial conditions outside and inside the limit cycle and integrate the trajectories until they reach a certain small Euclidean distance from the limit cycle, and then define our limit cycle strip as the space left. In this left over space we define quadrilaterals so as to fill up this ring. This becomes a strip in its own right, representing the limit cycle. Earlier we described the reversal mapping: mass reaching the end of a strip must be removed and deposited in a cell representing a stable point. Here, we use a similar approach: mass that arrives at the end of a strip must be removed and deposited on the limit cycle. We find the cell on the limit cycle that is closest in Euclidean distance to the limit cycle. Since the machinery to do this is already in place in the form of a reversal mapping, we will also refer here to this process as a reversal mapping. The modeler presents this reversal mapping in the same file format as used previously.

Initially we had attempted to define our limit cycle cells as having a fixed width, and then obtain strips by integrating backwards in time from the corners of these cells. Indeed, the coarse schematic grid in Fig. 14 has trajectories generated in this way for the interior of the limit cycle. However, for the purposes of actual computation, this method leads to degenerate cells. This is due to the fact that close to the limit cycle, trajectories move almost parallel to it, in particular along the “horizontal” segments of the limit cycle, where the fast *v* dynamics dominate. This leads to long, thin cells being created, which become degenerate when approaching the limit cycle - adjacent trajectories overlap to the degree of accuracy of the numerical integrator, leading to self-intersecting cells or cells with zero area.

From the outside of the limit cycle, most of the state space can be covered by simply choosing points on the edge of the region of interest and integrating forwards in time until one reaches the limit cycle. However, care must be taken when trajectories converge before arriving at the limit cycle, as shown in Fig. 14 B. This happens particularly along the cubic nullcline. We handle this by checking for degenerate cells or cells with area close to zero. These cells are then deleted from the grid, and instead a reversal mapping is created from the previous cell onto the closest (in Euclidean terms) cell.

The interior of the limit cycle proves to be even more challenging. Not only is there an unstable fixed point, also there exist canard trajectories, which have been the subject of considerable mathematical interest [28–31]. Loosely speaking, near the central portion of the cubic nullcline, there are slow but unstable trajectories. This leads to two types of numerical issues - first, the slow dynamics cause a build-up of exponentially many very small cells. We work around this by defining a minimum value of 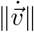 - regions below this value are considered to be approximately stationary, since they will have much slower dynamics than any noisy input we consider. The region we find is shown in Fig. 14 C.

We use cubic splines to approximate the boundary of this region, and then use points on this boundary as initial points for trajectories on the inside of the limit cycle to generate strips. Due to the instabilities in this region of the system, trajectories can be highly curved, and trajectories with initial conditions close to each other can diverge quickly, leading to cells which may intersect with each other, as shown in Fig. 14 E. As these areas with highly curved trajectories are still locally smooth, it may be possible to increase the resolution of the grid until non-degenerate cells are obtained, as we do here. However, it may not always be possible to do so due to computational constraints - in that case it may be more practical to delete bad cells after the creation of the grid and cover any gaps with fiducial bins.

To sum up, regions where trajectories merge - such as the limit cycle and nullclines in this case - involve moving from the two dimensional plane onto one dimensional trajectories, and pose conceptual as well as computational difficulties. Regions with highly curved trajectories may be possible to handle with very fine resolutions, but may pose difficulties at coarser resolutions. In both cases it is possible to handle such regions using an automated procedure: cells are checked for being complex quadrilaterals or having too small an area. Those satisfying this condition are deleted, and renewal mappings from the cells before them to the nearest cells are generated. Any gaps in the grid due to this can be handled using the prescribed method for creating fiducial bins.

In conclusion, we have successfully extended our procedure to dynamical systems with limit cycles and complex dynamics such as canards. While we have to make some compromises in the regions which pose significant analytic difficulty, these regions are those in which neurons would not spend any significant amount of time. Hence, our method would still be suitable for studying neural circuits of such populations.

### Numerical Solution and Efficiency

**Fig 15.**
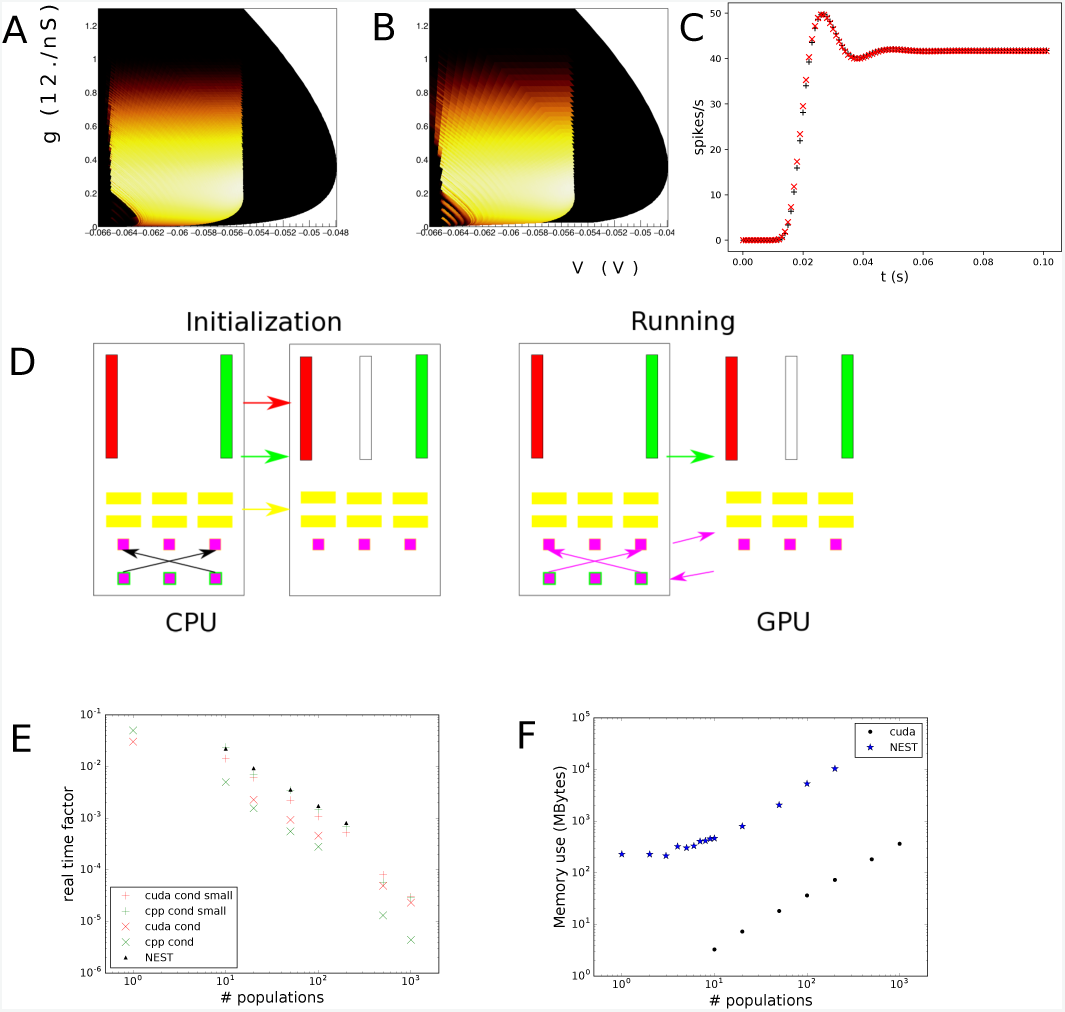
A-B: Comparison of the default mesh for conductance based neurons to a reduced mesh. Although coarser cells are visible in B, the density is not visibly affected, in particular in the white areas, where the bulk of the density is present. C: Firing rates are almost identical. D: Interaction between the C++ driver and the GPU registers, Interaction between CPU and GPU in a CUDA-based simulation. During the initialization, the mass array (red), the map array (green), the matrix elements (yellow) are copied onto the GPU. During simulation only the map (green) is updated in C++ and copied to the GPU, and firing rates are calculated from the meshes on the GPU. On the CPU the firing rates are processed by the network, and delays are applied where applicable. Resulting firing rates are sent back to the GPU for processing the the next simulation step. E: Run time factors for direct simulation, CUDA and C++ implementation (default and reduced mesh). F: memory use, NEST and CUDA implementation.

We solve Eq. 12 by a forward Euler scheme. Since we interleave moving probability mass through the grid with a numerical solution of Eq. 12, we solve Eq. 12 over a period Δ*t*, which can be as short as 10^−4^ s for some neural models. This renders sophisticated adaptive size solvers relatively inefficient. The matrices in Eq. 12 tend to be sparse band matrices, and one advantage of the forward Euler scheme is that it is embarrassingly parallel. For a single population in this we partition Δ*t* into *n*_*euler*_ time steps. For most simulations in this paper *n*_*euler*_ = 10 is adequate, we will discuss an exception below. In a forward Euler scheme Eq. 12 is discretized and a single step is given by:

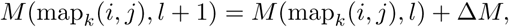

with

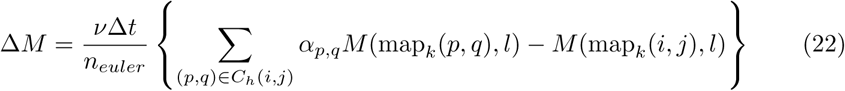

The current simulation time is *t*_*sim*_*k*Δ*t*, where Δ*t* is the mesh time step. The current map, which indicates where probability mass has moved under the influence of endogenous neural dynamics is therefore labelled by map_*k*_(*i*, *j*), which maps cell (*i,j*) to a unique mass array index as per Eq. 9. Note that the mapping should not be applied to the set *C*_*h*_(*i*, *j*). Simulation starts at *k* = 0, *l* = 0. Each Euler step l is increased, until *l* = *n*_*euler*_, upon which *l* is reset to 0 and *k* is increased by 1, until the desired end time is reached. The sets *C*_*h*_(*i*, *j*) and coefficients *α*_*p,q*_ remain constant throughout simulation.

Despite appearances, the right-hand side of Eq. 22 is of the form of a matrix vector multiplication, where the matrix is very sparse (Fig. 15 A). The matrix elements are numerical constants, and there is no dependence between rows, meaning that each row can be evaluated independently of the others and therefore the problem is extremely parallel: each row can be calculated in a separate thread when available.

Larger networks can be simulated by vectorizing the network: vectors representing the mass of individual populations are laid out in a single array representing the mass of multiple populations. Connections from population *i* to population *j* then are represented by block matrix elements, each consisting of one or more transition matrices (one for every efficacy associated with the connection) generated by the process described above.

We have created both a C++ and a CUDA implementation and evaluated them on single populations, as well as networks of populations. When generating networks we consider networks of conductance based populations, each population connected to constant input (‘cortical background’), with the network being sparsely connected (connectivity 5 %), efficacy chosen fixed (*h* = 0.05), with the number of connections drawn from a uniform distribution (*n* = 1 · · · 100). For small networks, the running times are not particularly onerous, with or without parallelization. For larger networks, in the C++ implementation we evaluated a single block matrix per thread using OpenMP. This means that in our implementation individual matrix vector calculations are not parallel, but that several matrix vector calculations are performed simultaneously. Since OpenMP offers a relatively small number of threads, this still makes efficient use of resources. The parallelization model for CUDA is different: we write a so-called kernel to evaluate Eq. 22 and launch a kernel for each block matrix. CUDA’s loop unrolling automatically performs parallelization within the kernel, and by launching kernels in different streams, inter kernel concurrency can be achieved. It is then the question whether the large number of threads compensate for the inherently slower GPU hardware. In Fig. 15 D we show how the GPU interacts with the C++ driver. During initialization the mass array is set up on the GPU, as well as the mapping, and the matrix elements. During a simulation run, the mass array mapping is updated, and firing rates are exchanged, but other than for visualization purposes, the mass array is not transferred, meaning lightweight communication between GPU and CPU.

We find that the number of cells in a mesh determines the performance. In Fig. 15 we examine the conductance based neuron example again. If we use the original mesh, without considering performance, we find that the method is slower than direct simulation as performed by NEST by a factor of three (6s per population for NEST −20s per population for the CUDA implementation). If we reduce the granularity of the original mesh, we find that we can bring the size of the mesh down from 120k cells to 25k cells, with the density and firing rate predictions unaffected (Fig.15 A vs B for density, B being the coarser mesh which is only visible in the bigger cells on top, C for firing rate). For the reduced mesh, the performance of the C++ implementation is equal to that of NEST, where the CUDA implementation is a little bit slower (measured on a Tesla P100). Both direct (NEST) simulation and C++ implementation use parallelization with 16 threads for this comparison. The real time factors (real time second divided by wall time of one simulated second) are shown in Fig. 15 E. A striking difference is the memory use (Fig. 15 F), which for the CUDA implementation is orders of magnitudes lower (300 MB for the largest network of 1000 populations, whereas a 100 population network with NEST already uses 10 GB). We conclude that the CUDA implementation supports the simulation of large networks on a single PC equipped with a GPGPU, whereas direct simulation requires a substantial cluster.

In the method we considered so far, a relatively large number of cells emerge around stationary points due to the exponential shrinkage of state space around them. In principle it is possible to group these cells together into larger ones, and to group them into strips that would run at a lower speed compared to the basic time step of the mesh. This would reduce the number of cells in the mesh considerably, while the basic granularity of the mesh will not be affected much. Such resulting merged cells are no longer quadrilateral and the method will have to be extended to be able to handle non-convex cells, which will be one of the first priorities in further work.

### General Remarks

We have demonstrated a very general method to study noise in 2D dynamical systems and applied them to various neural models and Tsodyks-Markram synapses. The state space of the deterministic model must be represented by a grid. The requirement that a grid be made is both a strength and a weakness: the state space relevant to the simulation must be chosen judiciously before the simulation starts. But since it must be constructed beforehand, integration can be done very accurately, using time steps that are much smaller than typically used in Monte Carlo simulators. If general purpose simulators are used with a default time step, and without adaptive methods that monitor errors, they may not alert the user to problematic regions of state space. Our method requires a careful layout of state space before simulation starts. We found that the requirement of a grid forces visualization and thereby already creates an understanding of the dynamics that can be expected.

When the state space cannot contain the simulation, this is clearly visible, either through loss of mass, or by the accumulation of mass at the edge of the grid. This proved useful in one instance, where a well known neural simulator produced a crash (due to an instability of the particular neural model implementation, not the simulator as such). Our method is very robust and stable, once a suitable grid is available. In general, we find that grids can be taken quite coarse in state space, but that a relatively small time step must be used for completely accurate results, such as comparison to analytic results like gain curves. When numerical errors are acceptable, and only qualitative agreement is required, much coarser grids can be used that require far less simulation time.

Our method is not as efficient as effective 1D methods [15–17], but makes very few assumptions. It handles time-dependent input without any restrictions. This is useful, for example, when comparing against basis functions expansions [13, 14,32]. These basis functions are typically determined for constant input, and time-dependent input must be treated as an adiabatic approximation. Our method does not require this. In short, our method may serve as benchmark for faster methods.

Population density techniques are part of an emerging ecology of simulation techniques, and it is important to consider their strength and weaknesses compared to related approaches. Direct spiking neuron simulations are straightforward in small to medium-sized networks, but hard to get right in large-scale simulations, where they are resource intensive (they can have large memory requirements, as well as being CPU intensive). The “missing spike” problem, and the need to keep track of spike information and its exchange between the various processors involved are just examples demonstrating that direct simulations are not straightforward. They have developed into a discipline of their own [36]. Population density techniques are conceptually simple, but unable to model pairwise correlations within a population, and the inclusion of finite-size effects is not straightforward (see [14] for an attempt). In general, population density techniques are not able to describe quenched network states, although fully connected networks are amenable to such analyses [37, 38]. Recently, a number of studies have explored path integral approaches to calculate pairwise correlations and to suggest a functional role for such correlations (e.g. [38–40]). Often, these techniques use the diffusion approximation, or are restricted to remain close to thermodynamic equilibrium. Two advantages of the technique described in this paper is that the latter restrictions do not apply. Theoretically, population density equations have been put on a rigorous mathematical footing [41], justifying its use for weakly connected networks where the quenched state of the network is not important. These papers also adds to a substantial body of observation that even for small networks population density techniques predict the firing rate correctly (e.g. [6,42] and many others). So, when modeling firing rates is the main objective, and the network is such that the populations may be far from equilibrium, population density techniques are a good candidate. They are also valuable in repeat experiments on single cells, as they show what noise will do to otherwise identical neurons.

Modeling a complex system of real neurons probably requires a hybrid approach. Mazzucato *et al.* [43] give an example of such an approach. They analyse the dimensionality of neural data recorded by multielectrode array. The dimensionality is estimated from pairwise correlations. They also produce a spiking network model, to validate their explanation and use population density techniques to establish the dynamical regime for their spiking network.

The study of 2D systems subject to noise is an important topic in its own right, given that limit cycles require at least two dimensions. The current trend in neuroscience towards 2D geometrical models reinforces this point.

An important prerequisite for the method to work is that the dynamical system can be represented faithfully. We found that some systems have challenging regions of state space: stationary points, whether stable or not, and limit cycles need careful handling and a full cover of state space is not possible. However, we find that we can infer motion of probability mass inside such regions from the immediate surroundings, the limit cycle of the Fitzhugh-Nagumo system as a case in point: it emerges as a region rather than as a curve from terminating the grid as it approaches the limit cycle.

There are interesting parallels between our method and a recently proposed method for determining missing spikes in hybrid time-driven, event-driven spiking neuron simulations [33]. Here, the authors consider the problem of missing spikes: the possibility that a neuron is below threshold at the end of a simulation step, but has crossed the threshold during the step. They solve this problem by determining whether a neuron is inside a volume in state space between the threshold and the backpropagated threshold. They find this easier than determining the actual point of crossing, and their method is reminiscent of ours when we calculate the transition matrix. They too consider a mapping like Eq. 6 which they are able to calculate explicitly for current based neurons. They conclude that apart from the threshold and the backpropagated threshold, the boundary is given by the vanishing tangent space of the map, precisely the criterion we used numerically (area of cell - in the absence of analytic solutions) to define boundaries of state space.

It is interesting to speculate about extending the method towards even higher dimensions. At first sight, this seems unfeasible: a three dimensional grid might already require millions of bins. It is not efficient to simulate systems with a size of the order 10^4^ particles by a larger number of bins. It would also be considerably harder to visualize the results. Nonetheless, probability tends to cluster in specific areas of state space and we find large parts of state space effectively unoccupied. A dynamical representation of the occupied part of state space would lead to a more scalable method. Our simulation results have shown that we can simulate large networks consisting of hundreds or thousands of populations. To make really large networks run more efficiently, we need smaller meshes and the best way to achieve that we now believe is to lump the large number of small cells that emerge near stationary points into larger ones, as described above. This will be our main focus for the near future.

## Acknowledgments

This project received funding from the European Union’s Horizon 2020 research and innovation programme under Grant Agreement No. 720270 (HBP SGA1) and Specific Grant Agreement No. 785907 (Human Brain Project SGA2). We thank Gaute Einevoll, Michele Giugliano and Viktor Jirsa for discussions, and Martín Pérez-Guevara for pushing large-scale use of the method.

